# Loss of CLN3 in microglia leads to impaired lipid metabolism and myelin turnover

**DOI:** 10.1101/2024.02.01.578018

**Authors:** Seda Yasa, Elisabeth S. Butz, Alessio Colombo, Uma Chandrachud, Luca Montore, Steven D. Sheridan, Stephan A. Müller, Stefan F. Lichtenthaler, Sabina Tahirovic, Susan L. Cotman

## Abstract

**Background:** Microglia are the primary brain cell type regulating neuroinflammation and they are important for healthy aging. Genes regulating microglial function are associated with an increased risk of neurodegenerative disease. Loss-of-function mutations in *CLN3*, which encodes an endolysosomal membrane protein, lead to the most common childhood-onset form of neurodegeneration, featuring early-stage neuroinflammation that long precedes neuronal cell loss. How loss of CLN3 function leads to this early neuroinflammation is not yet understood.

**Methods:** Here, we have comprehensively studied microglia from *Cln3*^Δex7/8^ mice, a genetically accurate CLN3 disease model. Microglia were isolated from young and old *Cln3*^Δex7/8^ mice for downstream molecular and functional studies.

**Results:** We show that loss of CLN3 function in microglia leads to classic age-dependent CLN3-disease lysosomal storage as well as an altered morphology of the lysosome, mitochonodria and Golgi compartments. Consistent with these morphological alterations, we also discovered pathological proteomic signatures implicating defects in lysosomal function and lipid metabolism processes at an early disease stage. CLN3-deficient microglia were unable to efficiently turnover myelin and metabolize its associated lipids, showing severe defects in lipid droplet formation and significant accumulation of cholesterol, phenotypes that were corrected by treatment with autophagy inducers and cholesterol lowering drugs. Finally, we observed reduced myelination in aging homozygous *Cln3*^Δex7/8^ mice suggesting altered myelin turnover by microglia impacts myelination in the CLN3-deficient brain.

**Conclusion:** Our results implicate a cell autonomous defect in CLN3-deficient microglia that impacts the ability of these cells to support neuronal cell health. These results strongly suggest microglial targeted therapies should be considered for CLN3 disease.

## Background

The neuronal ceroid lipofuscinoses (NCLs) are a group of rare inherited lysosomal disorders with overlapping clinical and pathological features, including seizures, progressive motor and cognitive decline and accumulation of autofluorescent material in the lysosome ^1^. The NCLs are caused by mutations in one of 13 distinct genes (*CLN1–CLN8*, *CLN10–CLN14)*, whose proteins have different characteristics, including subcellular localization and function ^1,2^. Despite the diversity of NCL proteins, their loss of function similarly causes aberrant lysosomal function, lysosomal accumulation of electron-dense autofluorescent storage material (primarily comprised of subunit c of the mitochondrial ATP synthase, saposins, and phospholipids), and cell death ^3,4^. NCL affects all ages and ethnicities, and the NCL subtypes are categorized with respect to the gene that causes the disorder ^5^. Autosomal recessive *CLN3* mutations cause CLN3 disease (also known as juvenile NCL or Batten disease), which is the most common form of neurodegeneration in children worldwide ^1^. Patients first develop vision loss, typically between 4 and 10 years of age, which is followed by onset of seizures, loss of motor and cognitive function, and premature death, typically by the mid-twenties ^3,4^. Unfortunately, there are no effective treatments for this devastating disorder.

Although the primary function of CLN3 is still unknown, numerous studies revealed that this endo/lysosomal transmembrane protein has roles associated with endocytic trafficking, autophagy, cytoskeletal organization, lysosomal pH homeostasis, lipid metabolism, and calcium signaling ^6–18^. While CLN3 disease is primarily a neurodegenerative disease, recent studies have demonstrated that peripheral organs are also clinically impacted at later disease stages ^19^. Despite the ubiquitous expression of *CLN3* ^20,21^, expression levels vary in different tissues and cell populations, suggesting the biological role of CLN3 may differ in different cell types ^20,22^.

Neuroinflammation has been documented at early disease stages in CLN3 disease mouse models. For example, astrocyte activation was observed already in 1–3-month-old homozygous *Cln3*^Δex7/8^ mice, whereas neuronal loss was not obvious until later ages (12–18 months) ^23,24^. Moreover, *in vitro* studies revealed the neurotoxic effects of CLN3-deficient primary microglia on neuronal growth in neuron-glia co-culture models ^25–27^. Constitutive activity of caspase-1 and concomitant inflammasome activation were also demonstrated in CLN3-deficient microglia, with increased levels of pro-inflammatory cytokines, including TNFα, IL-1β or IL-1α, following exposure to CLN3 disease-specific pathophysiological stimuli, such as ceramide or neuronal lysate ^26^. Taken together, these studies suggest neuroinflammation likely plays an important role early in the CLN3 disease process. Indeed, neuroinflammation has emerged as a candidate therapeutic target, with several recent studies investigating the impact of dampening glial activation on CLN3 neuropathogenesis with promising results ^25,27–29^. However, the mechanism by which neuroinflammation impacts disease onset and progression, especially how microglia play a role, is poorly understood. Establishing a clear role for loss of function mechanisms in CLN3-dependent neurodegeneration and a mechanistic connection to chronic inflammation has high potential to aid in developing novel therapeutic strategies in CLN3 disease. Here, we have established a model system to study CLN3-deficient microglia using acutely isolated microglia from homozygous *Cln3^Δex^*^7^*^/^*^8^ mice, which were engineered to recapitulate the most common CLN3 disease-associated mutation ^30^. Herein, we confirm a pro-inflammatory state and describe newly discovered molecular and functional changes in CLN3-deficient microglia that collectively point towards cell-autonomous defects in microglial function that likely impacts their ability to support neuronal cell health in CLN3 disease.

## Materials and methods

### Antibodies

The following antibodies were used: anti-Iba-1 [Western blot (WB): 1:1000, Wako 016-20001], anti-Iba-1 [immunofluorescence (IF): 1:500, Wako 019-19741], anti-Tuj-1 (WB: 1:1000, BioLegend MMS-435P), anti-GFAP (WB: 1:1000, Cell Signalling 3670S), anti-mCln3p (WB: 1:1000, ^10^), anti-Apoe (WB/IF: 1:250, Invitrogen 16H22L18), anti-PLP [WB: 1:1000, immunohistochemistry (IHC): 1:300, Abcam ab28486], anti-MBP (IHC: 1:1000, Sigma Aldrich SMI-99), anti-actin (WB: 1:1000, Santa Cruz Biotech sc-81178), anti-subunit c (IF: 1:300, Abcam ab181243 (ATP Synthase C [EPR13907]) and ^11^), anti-Lamp1 (WB: 1:1000, IF: 1:100, Santa Cruz Biotech sc-19992), anti-Lamp2a (WB: 1:1000, IF: 1:500, Abcam ab18528), anti-GM130 (IF: 1:100, BD Transduction 610822), anti-GRP175 (IF: 1:100, Stressgen SPS-825), anti-LRP1 (WB: 1:50000, Abcam ab92544), anti-Galectin 3 (WB: 1:500, Santa Cruz Biotech sc-32790), anti-PDI (IF: 1:100, StressGen SPA-891), secondary antibody goat anti-rat AlexaFluor 568 (1:800, Thermo Fisher Scientific Inc), secondary antibody donkey anti-rabbit AlexaFluor 488 (1:500, Thermo Fisher Scientific Inc) or 555 (1:800, Thermo Fisher Scientific Inc), secondary antibody donkey anti-mouse Alexa Fluor 488 (1:500, Thermo Fisher Scientific Inc) or 555 (1:800, Thermo Fisher Scientific Inc).

### Reagents

The following compounds were used in this study. ML-SA5 (10uM) (Cat no: AOB11116 (Aobious INC)), Verapamil (10uM) (Cat no: CAS 152-11-4 (Santa Cruz)), Lovastatin (10uM) (Cat no: S2061 (Selleck Chemicals)), LipiDyeII (1uM) (DiagnoCine FNK-FDV-0027), Filipin III (Sigma-Aldrich SAE0087), LysoTracker™ Deep Red (Invitrogen L12492), TrueBlack® Lipofuscin Autofluorescence Quencher (Biotium), ProLong™ Gold Antifade Mountant with DNA Stain DAPI (Invitrogen P36931), Tissue-Tek® O.C.T. Compound, ProLong™ Gold Antifade Mountant (P36930), PKH67 Green Fluorescent Cell Linker Kit for General Cell Membrane Labeling (Sigma-Aldrich Cat no: PKH67GL).

### Mice

*Cln3*^Δex7/8^ mice have been previously described and are publicly available at The Jackson Laboratory (Strain #: 017895) ^31^. *Cln3*^Δex7/8^ mice were maintained on a C57BL/6J congenic background (>20 generations) in the AAALAC-accredited rodent facility at MGH. Heterozygous matings were set up to generate wild-type, heterozygous, and homozygous littermate experimental animals, which were aged to the indicated timepoints for microglia isolation or tissues collection for histology. Mice were euthanized by inhalation of CO_2_-bottled gas, in accordance with current guidelines from the American Veterinary Medical Association. Mice collected for Histology studies were transcardially perfused, as described below. All procedures used for this study were reviewed and approved by the Subcommittee on Research Animal Care at MGH to ensure compliance with government regulations and animal welfare guidelines (Protocol # 2008N000013).

### Histology

Mice (N=3, 20-month old mice per genotype) were transcardially perfused with 4% PFA/0.1 M PBS, followed by brain dissection and six hours post-fixation with 4% PFA/0.1M PBS at 4°C. Brain hemispheres were placed into a scintillation vial and filled with 10%Sucrose/ 1X PBS. After 24 hours (24h), the sunken tissues at the bottom of the vial were transferred to 20%Sucrose/ 1X PBS. The next day, tissues were transferred to 30% sucrose/ 1X PBS. After 24h, the sunken tissues at the bottom of the vial were frozen in O.C.T. A Leica CM3050 S Cryostat was used to make 40 μm coronal brain sections collected in cryoprotectant solution (25% Ethylene Glycol/ 25% Glycerol/ 50% 1X PBS). Free-floating brain sections were processed for antigen retrieval in 1 mM sodium citrate buffer and microwaving for 10 minutes (min) at 10% power level. Brain sections were then blocked for 1h in 1X PBS/ 0.1% Triton X-100/ 1% Glycine/ 2% Bovine Serum Albumin (BSA)/ 10% Normal Horse Serum (NHS). Next, sections were incubated for two days at 4°C with primary antibody diluted in 1X PBS/ 2% BSA/ 10% NHS. After washing three times with 1X PBS/ 0.05% Tween 20, corresponding secondary antibodies were diluted in 1X PBS/ 2% BSA/ 10% NHS and incubated for 2h at room temperature (RT). Following secondary antibody incubation, sections were washed three times with 1X PBS/ 0.05% Tween 20 and incubated with Hoechst (1:10,000) for 10 min. Finally, sections were washed three more times in 1X PBS and subsequently mounted onto microscope slides in a bath of 1X PBS and allowed to dry overnight (O/N). Slides with the mounted sections were then coverslipped with mounting media and stored at 4°C in the dark until imaging. Imaging was performed on a Zeiss Celldiscoverer 7 microscope using a 20× / 0.95 Autocorr Objective.

### Primary microglia

Cerebellum, olfactory bulb, and brain stem were removed from dissected mouse brains. Meninges was dissected away from the remaining cerebrum in ice-cold HBSS under a dissection microscope. The cerebrum was then cut into 16 pieces and dissociated using the Neural Tissue Dissociation Kit (P) (Miltenyi Biotec), according to the manufacturer’s protocol. Microglial cells were isolated away from the rest of the cells in the suspension by incubation with CD11b microbeads (Miltenyi Biotec), which were then pulled down using a magnetic-activated cell sorting (MACS) separation LS column (Miltenyi Biotec). Samples prepared for mass spectrometry or biochemical analysis (e.g. immunoblotting) were further washed twice with PBS and frozen for subsequent analysis without culturing. Purity of the microglial fraction was confirmed by immunoblot with cell-type specific antibody markers (e.g. Iba1, microglia; Tuj1, neurons; GFAP, astrocytes). For cell culture experiments, isolated microglia were resuspended in microglial culture medium [DMEM/F12 (Gibco), 5% fetal bovine serum (FBS, Sigma Aldrich), 1% PenStrep (Gibco), and 10 ng/ml GM-CSF (R&D System)] and plated onto 12 mm glass coverslips. 50 µl, containing 8×10^4^ cells, was plated onto each coverslip placed inside of a well of a 24-well tissue culture plate. Cells were then left to attach for 30’ at 37°C, 5% CO_2_. After the 30-min incubation, 450 µl of culture media were added to each well containing the coverslips. The following day, media was exchanged for fresh media, and the cells were incubated for a further five days at 37°C, 5% CO_2_. After day five, cultured microglia were further processed for immunostaining, or labeling with LipiDyeII [1µM overnight (O/N) incubation at 37°C, 5% CO_2_] or Lysotracker (500nM for 30 min). For fixation, cells were washed three times with 1X PBS and fixed with 4% PFA for 15 min at RT. Alternatively, cells were fixed with ice-cold methanol: acetone (1:1) at −20°C for GRP75 and LAMP 1 immunostaining.

### Immunostaining

Fixed coverslips were further processed for immunostaining by first blocking and permeabilizing with 0.05% Triton X-100 in 10% NGS/PBS. To quench autofluorescence, coverslips were then treated with 1X TrueBlack solution for 30 seconds, followed by three washes with 1X PBS. Coverslips were then incubated with primary antibody diluted in 10% NGS/PBS O/N at 4°C. After the O/N incubation, coverslips were washed three times in 1X PBS, followed by a 1h RT incubation with secondary antibody diluted in 10% NGS/PBS (and where indicated, 100µg/ml Filipin dye for cholesterol staining was also added to the secondary diluted antibody solution). For LAMP 2A staining, 1% BSA/ 2% NHS/ 0.05% Saponin/ 1X PBS was used for blocking and antibody dilutions. Cells were washed and mounted with or without DAPI using Prolong Gold anti-fade mounting media, according to manufacturer guidelines. Samples were imaged on a Zeiss Epifluorescent Microscope equipped for digital capture of fluorescent images.

### Sample preparation for mass spectrometry

The microglial proteome of *Cln3^Δex^*^7/8/Δex7/8^ mice was compared with those of littermate control heterozygous (*Cln3^Δex^*^7/8/+^) or wildtype (*Cln3*^+/+^) mice at 3 (*Cln3^Δex^*^7/8/Δex7/8^ n=5 vs controls n=7) and 12 months (*Cln3^Δex^*^7/8/Δex7/8^ n=4 vs controls n=5) of age. Microglia were acutely isolated using MACS with CD11b microbeads (Miltenyi Biotec) as above and as previously described ^32^. Sample preparation and mass spectrometric measurements using data independent acquisition were performed as previously described ^33^. Data analysis was performed with Spectronaut 12.0.20491.14.21367 using a microglia spectral library generated in Monasor et al ^33^. Briefly, peptide and protein FDR were set to 1%. Protein label free quantification (LFQ) required at least two peptides. The statistical data analysis was performed in Perseus version 1.6.15.0 ^34^. Protein LFQ intensities were log2 transformed and two-sided Student’s T-tests were performed between *Cln3^Δex^*^7/8/Δex7/8^ and controls at 3 and 12 months to evaluate the significance of protein abundance changes. Only proteins with at least three valid values per group were considered for statistical testing. Additionally, a permutation-based FDR was used for multiple hypothesis correction ^35^ (p=0.05, s_0_=0.1).

The gene ontology enrichment for cellular component (GOTERM_CC_FAT) and biological process (GOTERM_BP_FAT) were carried out with the webtool DAVID ^36^. Proteins with an absolute log2 fold change larger than 0.5 and a p-value less than 0.05 were used as a target list and were compared with the list of all relatively quantified proteins as background.

### Immunoblot analysis

Acutely isolated microglia were lysed in lysis buffer (50mM Tris, 150mM NaCl, 0.2% Triton X-100) containing protease inhibitors (cOmplete Mini, Sigma Aldrich). Cells were completely lysed by passing the lysates through a 25G needle ten times followed by incubation on ice for 10 min. Lysates were then cleared by centrifugation at maximum speed in a Microfuge® 18 (Beckman) inside a 4°C cold room. Supernatants were collected and lysate protein content was quantified using a Pierce™ BCA Protein Assay Kit according to the manufacturer‘s protocol. 10-20 µg of total protein per sample was separated on a bis-tris acrylamide gel, followed by Western transfer and immunoblotting on nitrocellulose membrane (Bio-Rad). Immunoblots were developed using horseradish peroxidase-conjugated secondary antibodies (GE Healthcare) and the Western Lightning® Plus-ECL reagent (Perkin Elmer). Actin was used as a loading control. Developed immunoblots were imaged using an iBright FL1500 Imaging system (ThermoFisher). Immunoblot densitometry (n≥3 replicate samples) was performed using the gel analyzer tool in ImageJ (NIH).

### Crude myelin preparation and phagocytic assays

Dissected three-month-old wildtype mouse brains were sonicated in 10mM HEPES/ 5mM EDTA/ 0.3M Sucrose/ Protease inhibitors buffer. A sucrose gradient was prepared in ultracentrifuge tubes with 0.85M sucrose (pH 7.4) on the bottom and 0.32M sucrose (pH 7.4) on top. The brain homogenate was added to the sucrose gradient and centrifuged at 75,000 g for 30 min with a SW41 Ti rotor. The myelin fraction from the interface was collected in a new ultracentrifuge tube, and osmotic shock was applied with ultra-pure water by centrifuging the tubes at 75,000 g for 15 min three times. The yield of myelin was quantified by BCA assay, and some of the myelin was labelled with the green-fluorescent cell linker kit (PKH67 labeling kit, Sigma), according to manufacturer’s specifications.

For myelin phagocytosis, unlabeled or green fluorescent labeled myelin was resuspended with a 27G needle in microglial medium. Primary microglia were then incubated with 10 µg/ml of the resuspended myelin; for monitoring myelin uptake, cells were incubated with green fluorescent myelin for 2h in a 37 °C incubator ^32^, followed by three 1X PBS washes, and fixation with 4%PFA at RT for 15 min at the indicated timepoints. Myelin (green) phagocytosis inside phagocytic Lamp2a-positive vesicles (red) was monitored by fluorescent microscopy. For unlabeled myelin turnover studies, microglia were incubated with 10 µg/ml of the resuspended myelin for 6h at 37°C, 5% CO_2_. Myelin containing media was then removed, cells were washed three times with 1X PBS, and fresh microglial medium was added. Further incubation for the indicated times was followed by fixation and further processing for labeling and imaging studies. For microglia rescue experiments, after unlabeled myelin washout, microglia were incubated for an additional 48h, and for the last 24h, small molecule compounds (10µM concentration) were added (with or without LipiDyeII (1µM), as indicated). At the end of 48h, cells were fixed with 4% PFA at RT for 15 min and further processed for immunostaining.

### Transmission electron microscopy (TEM) imaging and analysis

Primary microglial cells were isolated as described above, the only exception being that the flow through from the MACS separation LS column was again loaded on the MACS separation MS column (Miltenyi Biotec) to ensure highest purity of the microglial fraction for TEM imaging. Brains from three 12-month-old mice/genotype were collected for three independent microglial preps/genotype for the TEM studies (N=3/genotype). The microglia pellets were collected and resuspended in 4% glutaraldehyde in cacodylate buffer. These pellets were resuspended in the buffer for 1hr at RT with gentle rotation. They were stored at 4°C until they were ready to be processed and sectioned for imaging.

Fixed isolated cell suspensions were centrifuged (3500rpm for 20min @ 4°C) to re-form a pellet aggregate, which was then resuspended for 1h at RT on a gentle rotator. Samples were rinsed multiple times in 0.1M cacodylate buffer, centrifuging to re-pellet as needed. Following supernatant removal, pellets were stabilized in 2% agarose in PBS, and the agarose-embedded pellets were then dehydrated using a graded series of ethanols to 100%, followed by a brief (10min) dehydration in 100% propylene oxide. Dehydrated samples were then pre-infiltrated O/N in a 1:1 mix of propylene oxide:Eponate resin (Ted Pella, Redding, CA) on a gentle rocker at RT. The following day, samples were placed into fresh 100% Eponate, which was allowed to infiltrate for several hours before embedding with additional 100% Eponate in flat molds. Resin was then allowed to polymerize 24-48h at 60°C. Thin (70nm) sections were cut on a Leica EM UC7 ultramicrotome using a diamond knife (Diatome U.S., Fort Washington, PA) and collected onto formvar-coated grids. Sections were then contrast-stained with 2% uranyl acetate and Reynold’s lead citrate, followed by examination on a JEOL JEM 1011 transmission electron microscope at 80 kV. Images were collected using an AMT digital camera and imaging system with proprietary image capture software (Advanced Microscopy Techniques, Danvers, MA).

For the purposes of morphometric quantification a minimum of 38 nucleated cells/ biological replicate sample were captured at 20,000X magnification. The number of storage bodies and lipid structures in each cell was then counted from intact cells in the images (i.e. full cell must have been visible in the image) until a total of 38 nucleated cells/biological replicate sample was reached (totaling 114 cells/genotype), and quantification data were further plotted and analyzed using GraphPad Prism.

### Motility assay

20,000 microglial cells were seeded per 96 Well Black/Clear Bottom Plate (TC Surface, cat number 165305). Microglial cells were imaged on a Incucyte® Live-Cell Analysis System (Sartorius), at 20X, every 15min, for 5h. Image sets were exported from the software as 8-bit .tif files, imported into FIJI image analysis software, and an image stack file was created. As described previously, the center of the nucleus on each image for every cell during the time course was tracked using the Manual Tracking plugin of ImageJ, and the velocity (µm/min) of each cell was quantified using the Chemotaxis Tool plugin of ImageJ and a Rose diagram graph data presentation ^37^.

### Image analysis

For intensity comparisons, color images were converted to gray, and using CellProfiler, both the primary (nucleus) and the secondary (cells) objects were identified. Each cell’s specific channel intensity was measured using the MeasureObjectIntensity module to analyze the total intensity of a specific channel in each cell.

For lysosomal perinuclear localization studies, the primary (nucleus; A), secondary (cell; B), and tertiary (lysosomes; B-A) objects were identified using CellProfiler. The nucleus was expanded twice by the ExpandOrShrinkObjects module to create a perinuclear region between the two layers that were created. Another tertiary object from that region was identified as perinuclear (C). The MeasureObjectInsensity module measured the perinuclear (C) and total (B-A) intensities of lysosomes. Then, perinuclear over total intensity ((B-A)/ C) was calculated for each cell.

For the Golgi compactness calculation, the primary (nucleus; A), secondary (cell; B), and tertiary (Golgi; B-A) objects were identified using CellProfiler. The MeasureObjectSizeShape module was used to see each cell’s compactness value for the Golgi.

For the mitochondrial fragmentation and length analysis, the primary (nucleus; A), secondary (cell; B), and tertiary (mitochondria; B-A) objects were identified using CellProfiler. Mitochondria features were suppressed and thresholded, and the MorphologicalSkeleton module was used to get the mitochondrial skeleton. Primary objects were identified from the skeleton image to identify each mitochondria in each cell. Mitochondrial primary objects were shrunk, and endpoints were added by the Morph module. Primary objects were identified and shrunk from the endpoint data. Lastly, the MeasureObjectskeleton module was used to have the values for mitochondrial fragmentation and the length of each mitochondria from every cell ^38^.

For the LD analysis, primary objects were identified and expanded to identify cells using CellProfiler. The IdentifyPrimaryObjects module was used again to identify LDs from the identified cells’ images. The RelateObjects module was used to identify the LDs of each cell and calculate their numbers in every single cell.

For MBP and PLP fluorescent intensity measurement, tile scan images captured using the Zeiss Celldiscoverer 7 microscope were initially processed through the ImageJ “Make Binary” tool for binary image conversion. Subsequently, the region of interest was delineated using the ImageJ “Freehand Selections” tool on the binary image. Region of interest was the outline of the tissue section, with folds in the tissue excluded. The ImageJ “Measure” tool was employed to directly assess the mean intensity values for MBP and PLP fluorescence by utilizing the “mean grey value” readout. Mean MBP and PLP fluorescent intensities were measured in three separate brain sections per animal, which were then averaged for subsequent statistical analysis, as done previously ^39^.

### Statistical analysis

An unpaired Student’s t-test with a two-tailed distribution was used to compare the two groups **(Supplementary Figures 3, 4) (Fig. 1, 2, 3, 4c**, 6). One-way ANOVA followed by Tukey’s multiple comparisons test was used to compare multiple groups **(Fig. 4n)**. Two-way ANOVA followed by Sidac’s multiple comparison test was used to compare the mean differences between groups split into two independent variables **(Fig. 5)**. GraphPad Prism 10.0.2 was used for all. A value of P ≤ 0.05 was considered significant (*P ≤ 0.05; **P ≤ 0.01; ***P ≤ 0.001; ****P ≤ 0.0001).

**Fig. 1.**
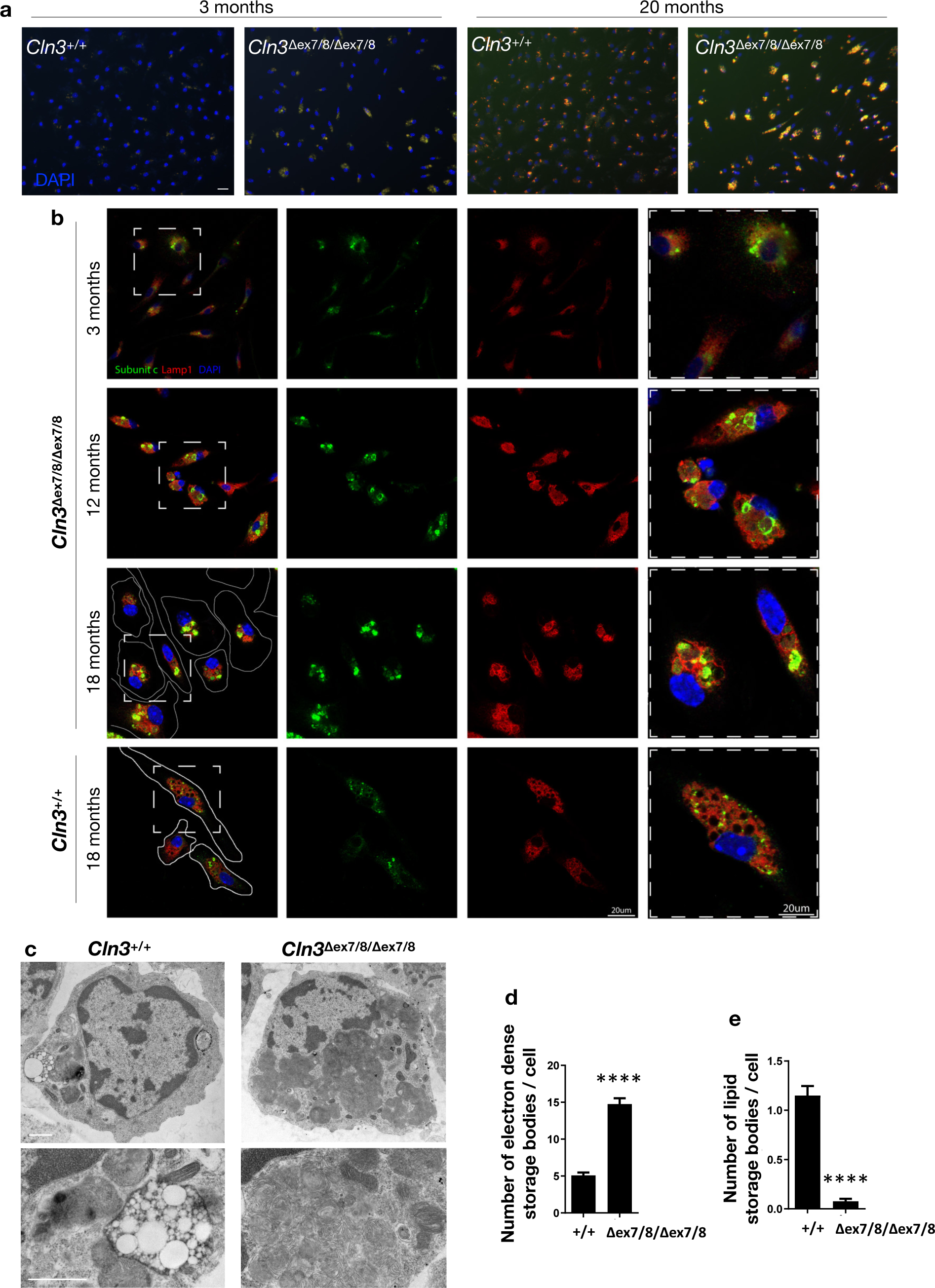
CLN3-deficient microglia accumulate high amounts of storage material. **a)** Representative fluorescent images of cultured 3- and 20-month-old (mo) microglia from *Cln3*Δex7/8/Δex7/8 mice and wild-type (*Cln3*+/+) littermates. Microglia from aged mice show broad-spectrum autofluorescence, which is more abundant in the *Cln3*Δex7/8/Δex7/8 microglia. Scale bar= 25 µm. **b)** Representative micrographs of cultured primary isolated microglia from 3-, 12-, and 18-month-old homozygous *Cln3*Δex7/8 mice and from 18-month-old wild-type littermate mice for comparison. Microglia were immunostained with Lamp1 (red) and subunit c antibodies (green). Scale bar= 20 µm. **c)** Transmission electron microscopy (TEM) images of acutely isolated microglia from *Cln3*Δex7/8/Δex7/8 mice and littermate wild-type controls at the age of 12 months. Scale bars= 800 nm. **d,e)** Bar graphs showing the quantification of storage material and lipid storage bodies from TEM images. N=114 cells/genotype, from 3 mice per genotype (38 cells/mouse). Student’s t-test, ****P ≤ 0.0001.

**Fig. 2.**
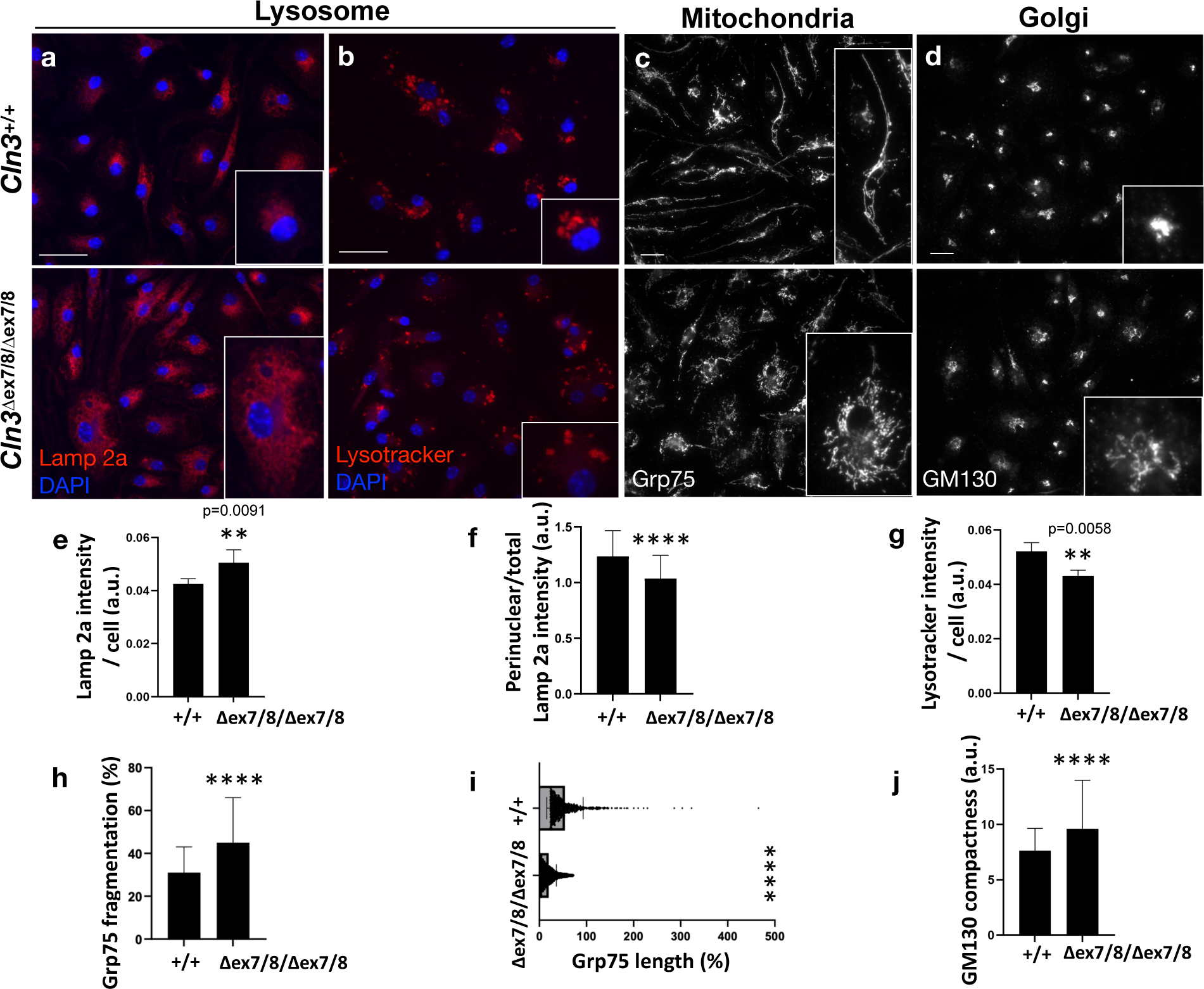
Altered organellar morphology in CLN3-deficient microglia. Cultured microglia from 6-month-old mice were stained with lysosomal markers, Lamp2a (red) **(a)** or LysotrackerTM **(b)**, mitochondrial marker, Grp75 (white) **(c),** and Golgi marker, GM130 (white) **(d).** Quantification of morphological changes is shown in the bar graphs **(e-j)**. GM130 compactness determination using CellProfiler **(j)** assigns circular objects a compactness value of 1 and irregular objects, or objects with holes, compactness values greater than 1, such that a higher value indicates reduced compactness. N= 300 cells/ genotype (3 mice/genotype from 3 independent experiments and 100 cells/mouse). Unpaired Student’s t-test with a two-tailed distribution was used to compare the two groups. Scale bars= 25 µm. Data are shown as mean ± s.d.; **P ≤ 0.01, ****P ≤ 0.0001 Student’s t-test.

**Fig. 3.**
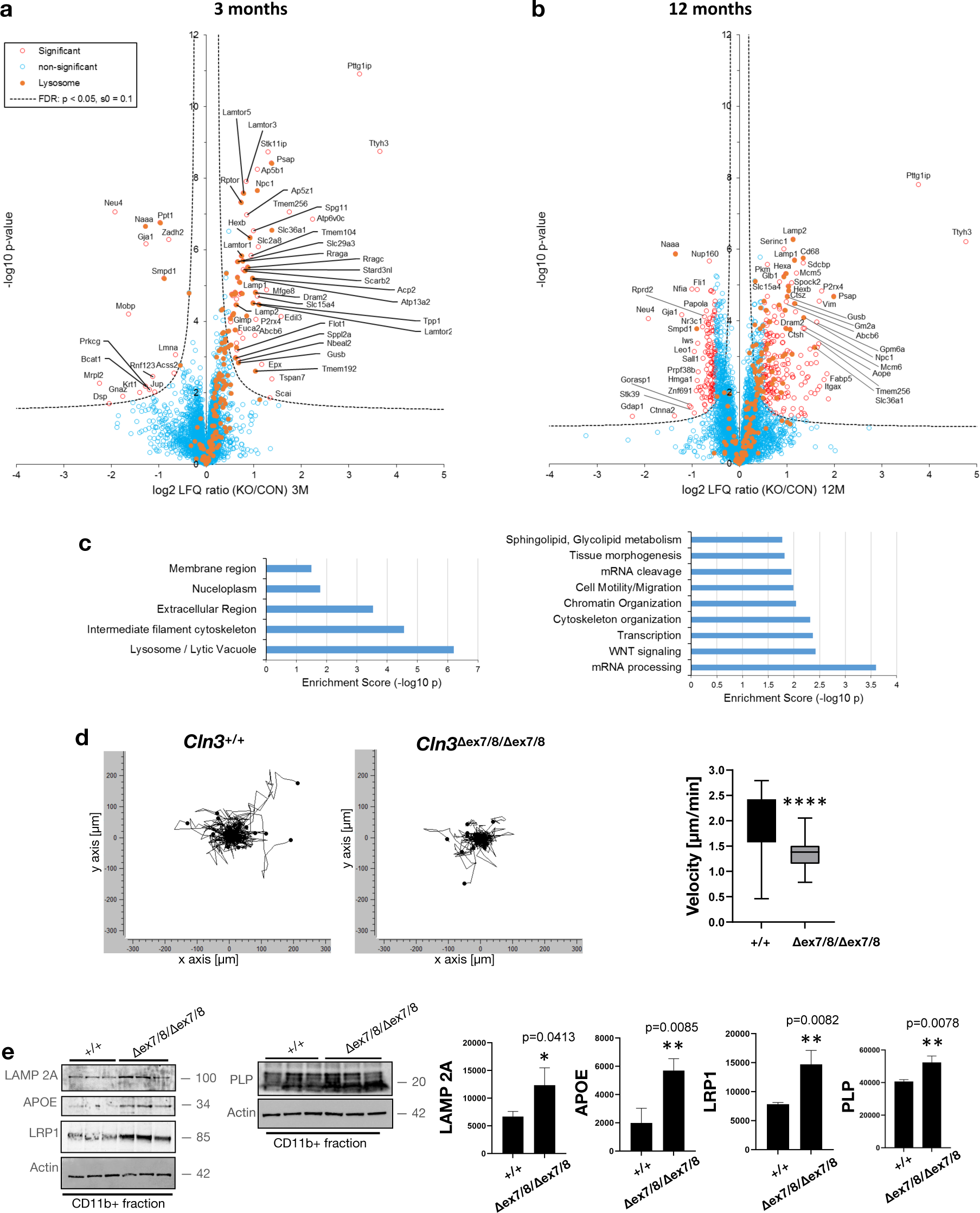
Proteomic analysis of homozygous *Cln3*Δex7/8 microglia reveals early lysosomal changes. Relative levels of detected proteins for homozygous *Cln3*Δex7/8 microglia (KO) compared to littermate heterozygous and wild-type control microglia (CON) are depicted in the volcano plots from 3-month **(a)** and 12-month-old mice **(b)**. Open red dots indicate proteins with a significant change in abundance [greater than false discovery rate (FDR, hyperbolic curve) threshold], and orange dots represent lysosomal proteins, while open blue dots represent proteins that were detected but did not significantly differ between KO and controls. **(c)** GO analysis results for ‘Cellular component’ (left) and ‘Biological process’ (right). **d)** Graphs represent results of a representative migration assay on microglia isolated from homozygous *Cln3*Δex7/8 and wild-type control mice. Real-time imaging was performed every 15 minutes for 5 hours to generate the migration plots, with each track from an individual cell aligned at the starting point. Axis units in µm from the starting point. Quantification of migration velocity (µm/min) over 5 hours was performed using ImageJ and Ibidi’s Chemotaxis and Migration Tool plugin. N= 37 cells for wild-type and N= 45 cells or homozygous *Cln3*Δex7/8 microglia. Data are shown as mean ± s.d.; ****P ≤ 0.0001 Student’s t-test. **e)** Bar graphs show the quantification of immunoblots using ImageJ (N= 3 biological replicate samples/genotype). Data are shown as mean ± s.d.; *P ≤ 0.05, **P ≤ 0.01 Student’s t-test.

**Fig. 4.**
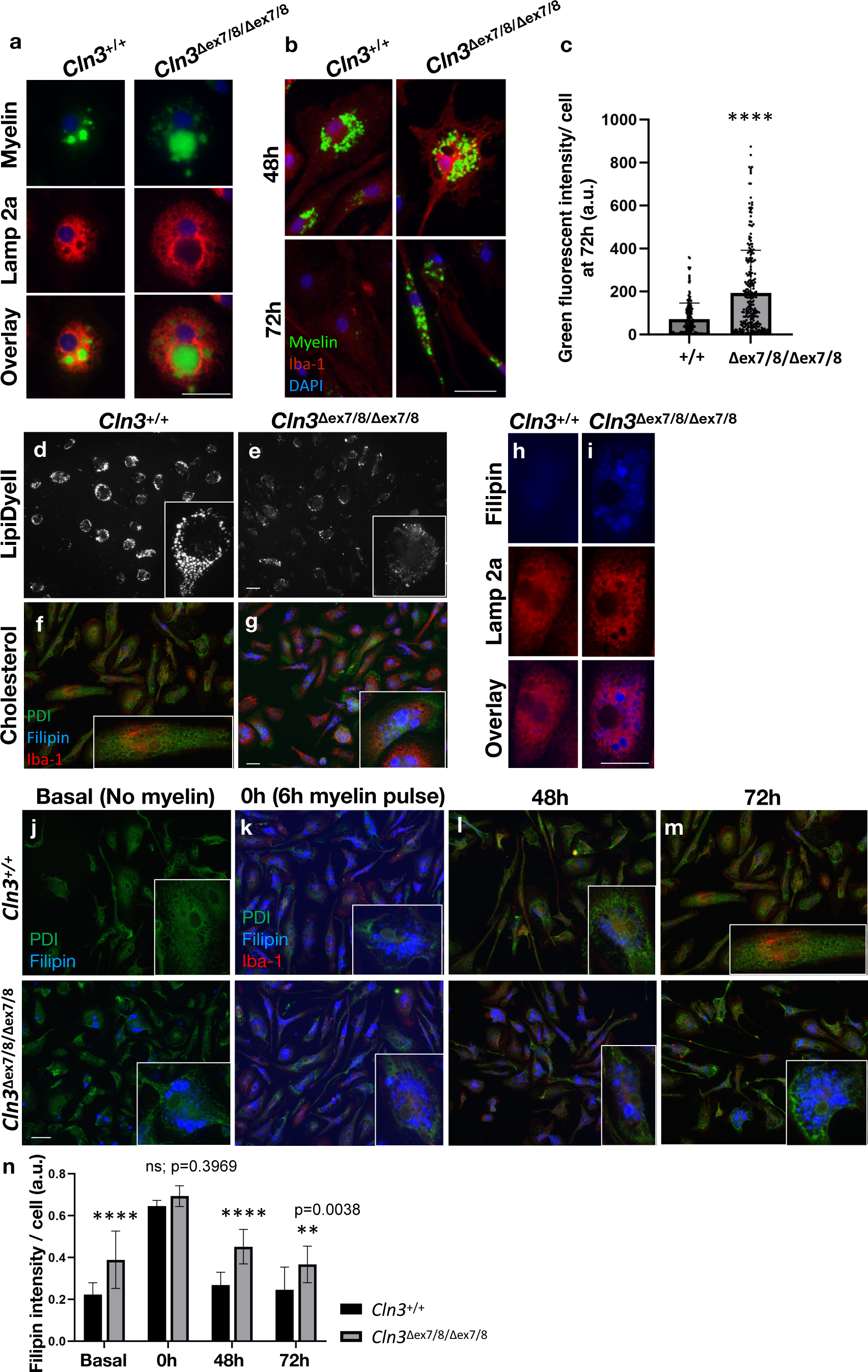
Myelin turnover is disrupted in CLN3-deficient microglia. **a)** Representative micrographs of green-fluorescent myelin uptake for wild-type and homozygous *Cln3*Δex7/8 microglia. Myelin (green), Lamp2 (red), DAPI= nuclei (blue). See Supplementary Figure 4A for low-power images. **b)** Representative micrographs of green-fluorescent myelin degradation at 48 and 72 hours for wild-type and homozygous *Cln3*Δex7/8 microglia. Myelin (green), Iba-1 (red), DAPI= nuclei (blue). See Supplementary Figure 4B for low-power images. **c)** Quantification of green fluorescent-myelin at 72 hours (N= 203 cells/wild-type, N= 228 cells/*Cln3*Δex7/8/Δex7/8 microglia, from 3 independent experiments). Each dot represents one cell. Data are shown as mean ± s.d.; ****P ≤ 0.0001, Student’s t-test. See Supplementary Figure 4B for quantification of green fluorescent-myelin at 48 hours. **d-g)** Representative micrographs of unlabeled myelin uptake assays followed by LipiDyeII labeling of lipid droplets and Filipin labeling of cholesterol, 48 hours post myelin washout. **h, i)** Zoomed in representative images from unchallenged (i.e. no myelin phagocytosis) 6-month-old wild-type and *Cln3*Δex7/8 mice microglia co-stained with Filipin (blue) and lysosomal marker Lamp2a (red). See Supplementary Figure 4C for low-power images. **(j-m)** Representative micrographs following Filipin staining during myelin uptake and turnover. Basal panels show Filipin staining prior to myelin uptake. PDI (green), Iba-1 (red), and Filipin (blue). **n)** Bar graph shows quantification of Filipin intensity per cell. (N= 210 cells/wild-type and N= 230 cells/*Cln3*Δex7/8/Δex7/8 from 3 independent experiments) Data are shown as mean ± s.d.; ns: non-significant, **P ≤ 0.01, ****P ≤ 0.0001, Two-way ANOVA followed by Sidac’s multiple comparison test. Scale bars= 25 µm.

**Fig. 5.**
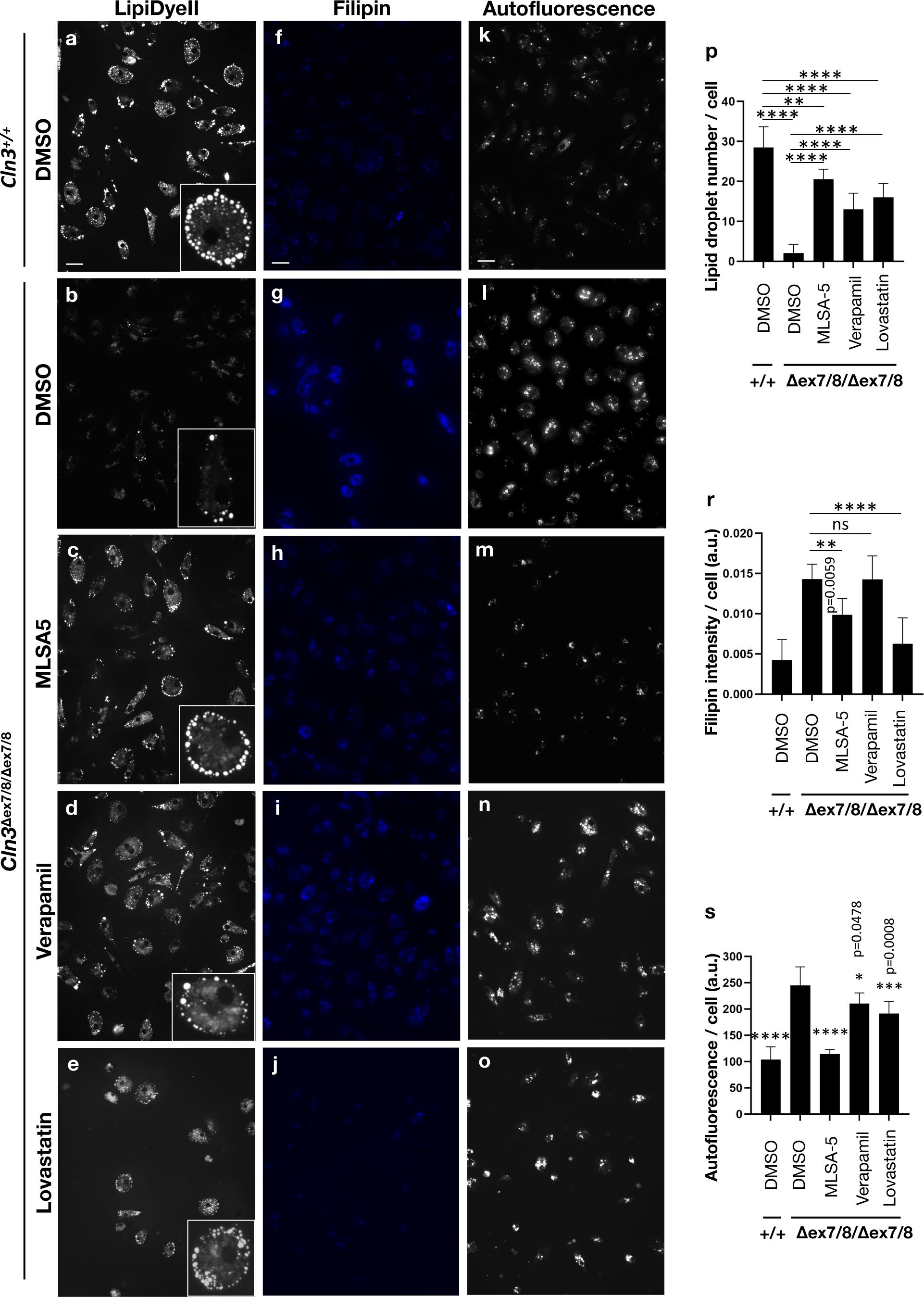
Correction of storage material and myelin turnover defects in CLN3-deficient microglia using small molecules targeting autophagy/ lysosomal function and cholesterol. **(a-j)** Representative micrographs of wild-type and *Cln3*Δex7/8/Δex7/8 microglia, treated with small molecules or DMSO vehicle, as indicated, and subsequently labeled with LipiDyeII or Filipin, 48 hours post-myelin washout. For autofluorescence, microglia were unchallenged (no myelin) and unlabeled. Scale bars= 25 µm. **(p)** Bar graph shows quantification of lipid droplet number (N=200 cells/genotype or treatment, from 3 independent experiments). **(r)** Bar graph shows quantification of Filipin intensity (N= 240 cells/genotype or treatment, from 3 independent experiments). **(s)** Bar graphs show quantification of autofluorescence intensity (N= 180 cells/genotype or treatment, from 3 independent experiments). Data are shown as mean ± s.d.; ns: non-significant, *P ≤ 0.05, **P ≤ 0.01, ***P ≤ 0.001, ****P ≤ 0.0001, One-way ANOVA followed by Tukey’s multiple comparisons test was used.

**Fig. 6.**
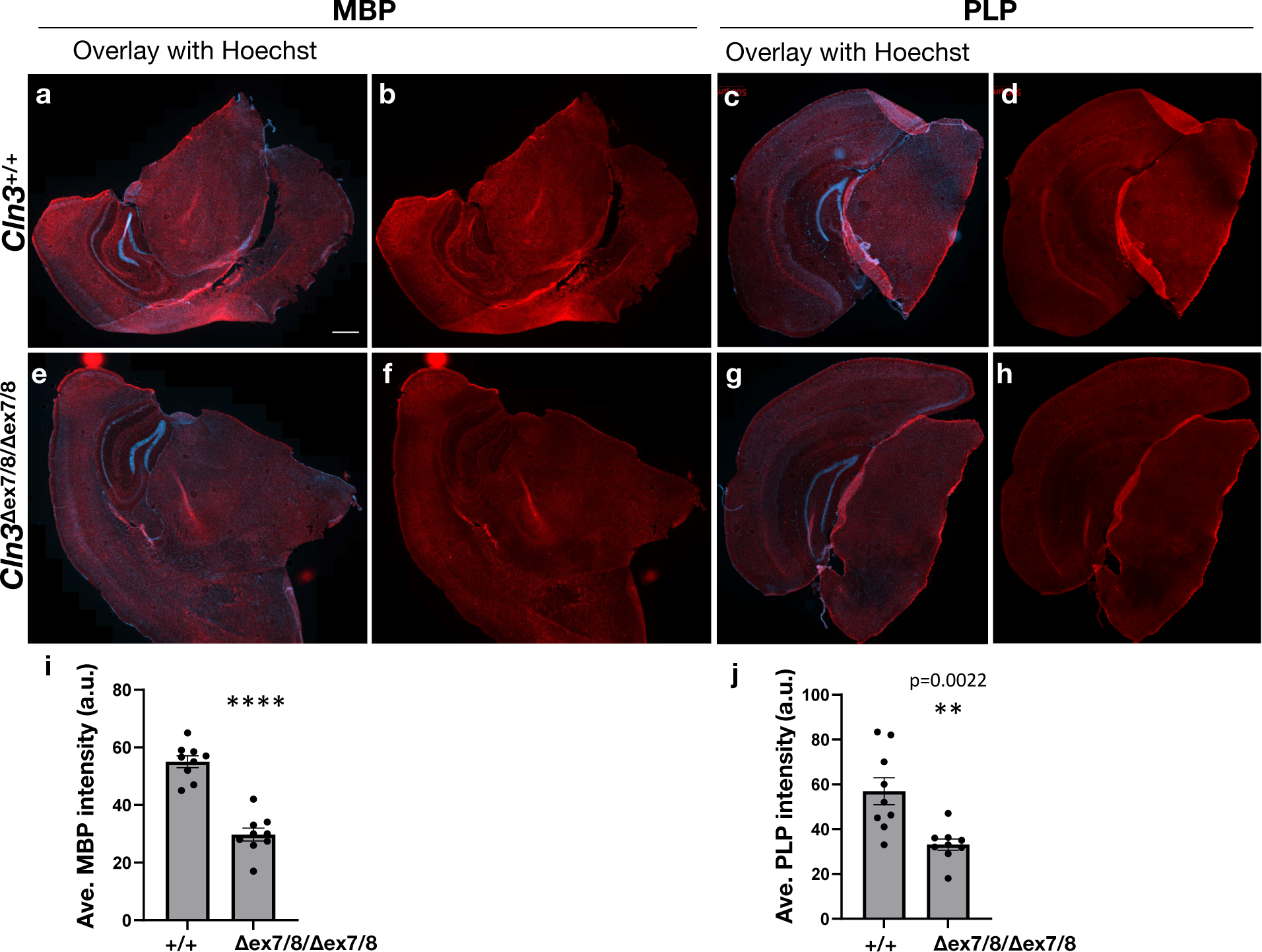
Reduced myelin staining in aged *Cln3*Δex7/8/Δex7/8 mouse brain. Representative brain sections from 20-month-old wild-type **(a-d)** and *Cln3*Δex7/8/Δex7/8 **(e-h)** mice immunostained for myelin using anti-MBP (red) or anti-PLP (red) antibody. Hoechst nuclear counterstaining is shown in blue. Bar graphs represent the average of mean intensities **(i,j)**. N=9 brain sections (3 mice per genotype, 3 brain sections per mouse, matched to represent comparable brain regions for control and *Cln3*Δex7/8/Δex7/8). Data are represented as mean ± standard error of the mean (SEM); **P ≤ 0.01, ****P ≤ 0.0001, Student’s t-test. Scale bar= 500 µm.

## Results

### *Cln3*^Δex7/8/Δex7/8^ mouse microglia exhibit characteristic pathological features of CLN3 disease

To study CLN3-deficient microglia, we used a MACS approach to establish highly purified microglial fractions from adult homozygous *Cln3*^Δex7/8^ mouse brain for downstream primary culture and morphological and biochemical analysis ^32^ (**Supplementary Figure 1a**). The hallmark pathology of CLN3 disease is electron-dense autofluorescent ceroid lipopigment accumulation inside lysosomes ^1^. Thus, we first isolated microglia from homozygous *Cln3*^Δex7/8^ mice and wild-type littermates at timepoints between 3 and 20 months-of-age, which were then evaluated for the relative burden of storage material. As early as 3 months, isolated microglia from homozygous *Cln3*^Δex7/8^ mice had significantly more autofluorescent storage material than that observed in wild-type microglia **(Fig. 1a**; yellow signal). With increasing age, the amount of autofluorescent material increased in both wild-type and homozygous *Cln3*^Δex7/8^ microglia, but this was more dramatic in the homozygous *Cln3*^Δex7/8^ microglia **(Fig. 1a)**. Subunit c of the mitochondrial ATP synthase (hereafter referred to as “subunit c”) is the main accumulated protein inside CLN3-deficient lysosomes ^3,4^, and indeed, the storage material in homozygous *Cln3*^Δex7/8^ mouse microglia was immunopositive for subunit c. As shown in Figure 1b, the relative burden of subunit c-positive storage material substantially increased with age, such that by 18 months-of-age, large subunit c-positive deposits were detected in swollen Lamp1-positive vesicles. These findings were consistent with a previous study in which a subpopulation of microglia was shown to accumulate autofluorescent lysosomal storage material in aging wild-type mice, and this population of microglia from homozygous *Cln3*^Δex7/8^ mice accumulated dramatically more autofluorescent material compared to that observed in age-matched wild-type littermate microglia ^40^.

To examine the ultrastructure of the lysosomal storage material in the homozygous *Cln3*^Δex7/8^ mouse microglia, we also performed TEM on isolated microglia from 12-month-old mice. As shown in Figure 1, the cytoplasm of the homozygous *Cln3*^Δex7/8^ microglia was filled with electron-dense, membranous storage bodies, which at higher power magnification exhibited a multilamellar fingerprint-like profile, consistent with the classical NCL-type storage material that is seen in CLN3 disease **(Fig. 1c,d)** ^1^. By comparison, only occasional membranous structures were observed in the cytoplasm of wild-type microglia, which also frequently contained electron-lucent lipid storage bodies similar to those previously described in aged autofluorescent-positive microglia ^40^. However, interestingly, these lipid storage bodies were virtually absent in the microglia from homozygous *Cln3*^Δex7/8^ mice **(Fig. 1e)**.

It has been shown in neurons and in lower organism models that, in addition to endolysosomal abnormalities, other organelle systems are also impacted by the loss of CLN3 function ^6,15,41,42^. However, to the best of our knowledge, this has not been examined in microglia. Therefore, we next wanted to examine late endosomes/lysosomes more carefully and to survey the membrane organelles in microglia isolated from homozygous *Cln3*^Δex7/8^ mice. To accomplish this, we probed the isolated microglia from homozygous *Cln3*^Δex7/8^ and wild-type littermate mice with additional markers for endolysosomes, endoplasmic reticulum, Golgi apparatus, and mitochondria (lysosome-associated membrane protein 2a ‘Lamp 2a’, LysoTracker™, PDI, GM130, Grp75, respectively). Consistent with our observations using Lamp 1 antibody **(Fig. 1b)**, Lamp 2a immunostaining highlighted a significantly expanded lysosomal network in homozygous *Cln3*^Δex7/8^ microglia compared to wild-type controls, where both the Lamp 2a signal intensity and the relative distribution of the Lamp 2a labeled structures differed (**Fig. 2a,e,f**). Further staining using LysoTracker™, a pH-sensitive lysosomotropic dye, also supported these observations, resulting in a staining pattern of more peripherally distributed lysosomes in the homozygous *Cln3*^Δex7/8^ microglia compared to wild-type controls **(Fig. 2b)**. Notably, the relative intensity of LysoTracker™ signal was also reduced in the homozygous *Cln3*^Δex7/8^ microglia compared to wild-type **(Fig. 2g)**, suggesting CLN3-deficient microglia may have an altered lysosomal pH, as reported in several other cell models of CLN3 disease ^14–16^.

In addition to the lysosomal abnormalities observed in homozygous *Cln3*^Δex7/8^ microglia, we also observed altered morphology of the mitochondrial network and the Golgi apparatus **(Fig. 2c,d,h-j)**. More specifically, the mitochondrial network appeared more fragmented in homozygous *Cln3*^Δex7/8^ microglia compared to wild-type **(Fig. 2h,i)**, and the Golgi was less compact and more spread away from the nucleus in the homozygous *Cln3*^Δex7/8^ microglia compared to wild-type **(Fig. 2j)**. No obvious morphological differences were observed in the ER, as highlighted by PDI immunostaining **(Supplementary Figure 2)**.

### *Cln3*^Δex7/8/Δex7/8^ mouse microglia harbor a significantly altered lysosomal proteome

To gain further unbiased insights into the impact of CLN3 deficiency on microglia, we prepared acutely isolated microglia samples from 3-month-old and 12-month-old homozygous *Cln3*^Δex7/8/Δex7/8^ mice and littermate controls (wild-type or heterozygous littermates), which were then subjected to proteomics analysis by liquid chromatography/mass spectrometry (LC/MS). Publicly available RNA sequencing data indicated that the *Cln3* expression level in mouse brain is relatively higher in microglia compared to other cell types, including neurons, oligodendrocytes, and astrocytes ^43^. Notably, CLN3-specific peptides were detected by LC/MS in both 3-month-old and 12-month-old control microglial samples, and CLN3 was also detected by immunoblot analysis in microglia-enriched samples from wild-type mice, more than in the microglia-depleted fractions from wild-type mice containing primarily neurons and astrocytes, consistent with the RNAseq data **(Supplementary Table 1; Supplementary Figure 1)**. To the contrary, no CLN3 peptides were detected by LC/MS in the homozygous *Cln3*^Δex7/8/Δex7/8^ microglial samples **(Supplementary Table 1)**.

As depicted by the volcano plots in Figure 3, our proteomics analysis revealed prominent upregulation of lysosomal proteins already at 3 months-of-age **(Fig. 3a)**, and this was exacerbated further with age, with both a greater degree of change and a greater number of lysosomal proteins with increased abundance at 12 months-of-age in homozygous *Cln3*^Δex7/8^ microglia compared to control microglia **(Fig. 3b)**. Moreover, multiple components of the mTOR pathway were also upregulated at both 3 and 12 months-of-age **(Fig. 3a,b)**, including the mTORC1 signaling complex (mTOR, Rptor, Lamtor1, 2, 3, and 5, and Mlst8); mTORC1 is a major signaling hub associated with the lysosome and metabolic regulation ^44^. The two most dramatically changed proteins, which were both increased ∼10-fold or more in homozygous *Cln3*^Δex7/8^ microglia compared to control microglia at both the 3- and 12-month timepoints, were relatively uncharacterized lysosomal membrane proteins, PTTG1IP and TTYH3 **(Fig. 3, Supplementary Table 1)** ^45^.

We next performed a Gene Ontology (GO) analysis on the 12 months-of-age proteomic datasets to gain more comprehensive insights into the pathways affected by loss of CLN3 function in microglia. Not surprisingly, the lysosome/lytic vacuole was the most prominently changed cellular component, with the intermediate filament cytoskeleton next most prominent **(Fig. 3c)**. Among the lysosomal proteins upregulated were Lamp 1 and Lamp 2, consistent with our immunostaining studies and prior reports **(Fig. 2**, ^40^), and further validated by immunoblot **(Supplementary Figure 3a, Fig. 3e)**. GO biological processes that were significantly altered in homozygous *Cln3*^Δex7/8^ microglia compared to wild-type were mRNA processing, WNT Signaling, Transcription, Cytoskeleton Organization, Chromatin Organization, Cell Motility/ Migration, mRNA Cleavage, Tissue Morphogenesis, and Sphingolipid, Glycolipid Metabolism, revealing numerous interesting pathways for further study **(Fig. 3c)**. Given the GO analysis results highlighting cytoskeleton and cell motility/migration, combined with multiple prior studies showing cytoskeleton and motility defects in other cell types with CLN3-deficiency ^46,47^, we tested the motility of the cultured microglia isolated from homozygous *Cln3*^Δex7/8^ and wild-type mice using live cell time-lapse imaging. As shown in Figure 3, *Cln3*^Δex7/8/Δex7/8^ microglia showed significantly reduced motility over a 5-hour period, as compared to wild-type microglia **(Fig. 3d)**.

In addition to the GO biological process analysis revealing enrichment in the “Sphingolipid, Glycolipid Metabolism” pathway, we also noted a number of individual lipid-related proteins with significantly altered levels in both the early and later timepoints (3 and 12 months-of-age, respectively). Notable examples include sphingomyelin phosphodiesterase 1 (SMPD1), proteins linked to glutathione metabolism [glutathione S-transferase omega 1 (GSTO1)], myelin-associated oligodendrocyte basic protein (MOBP), prosaposin (PSAP), and low-density lipoprotein receptor-related protein 1 (LRP1), which is important in lipoprotein metabolism and cell motility ^48^. Finally, consistent with prior reports, homozygous *Cln3*^Δex7/8^ microglia showed elevated levels of multiple inflammatory markers, [apolipoprotein E (APOE), integrin alpha-X (ITGAX), CD68, vimentin, galectin-3] **(Supplementary Table 1, Supplementary Figure 3a,b, Fig. 3e)**. Notably, however, these were less prominent changes than what was seen for lysosomal proteins and mostly these were only significantly upregulated at the 12-month timepoint. The altered levels for several of these proteins were validated in additional microglial samples, including APOE and LRP1 **(Fig. 3e)**.

### Abnormal myelin turnover and cholesterol accumulation in *Cln3*^Δex7/8/Δex7/8^ microglia

Myelin, an important insulating component of neurons that is important for transduction of electrical impulses along neuronal axons, can be taken up by microglia via phagocytosis for turnover, where it is trafficked to the lysosome for degradation by proteases and lysosomal lipases. The cholesterol component of myelin cannot be degraded by lysosomal enzymes and instead is exported from the lysosome for extracellular release, or it is transported to the ER where it is esterified and stored in lipid droplets (LD), as cholesterol esters. Cholesterol esters can then be used by the cell as energy fuel ^49^. These pathways for handling free cholesterol are critical for cellular health since accumulated cholesterol is toxic to the cell ^50^. In addition to the altered levels of lysosome and lipid-related proteins in CLN3-deficient microglia detected through our proteomics studies, we also noted altered levels of several myelin-component and myelin-related proteins in the *Cln3*^Δex7/8/Δex7/8^ mouse microglial proteome. For example, myelin proteolipid protein 1 (PLP), myelin basic protein (MBP), and LRP1 (an essential receptor for microglial myelin phagocytosis ^51^) all significantly differed in the *Cln3*^Δex7/8/Δex7/8^ microglia, compared to control littermate microglia **(Supplementary Table 1)**. Interestingly, PLP levels were significantly lower at 3 months-of-age, but they were significantly elevated in microglia isolated from 12-month-old *Cln3*^Δex7/8/Δex7/8^ mice **(Supplementary Table 1, Fig. 3e)** compared to control littermates. Given these observations, we next evaluated myelin phagocytosis and LD formation in CLN3-deficient microglia to test whether the proteomic signatures observed were accompanied by functional changes.

To first study myelin uptake and degradation, we incubated green fluorescent-labeled myelin with microglia isolated from wild-type and *Cln3*^Δex7/8/Δex7/8^ mice. We did not observe an obvious difference in relative green fluorescent-myelin uptake between *Cln3*^Δex7/8/Δex7/8^ and wild-type microglia at the 2-hour timepoint, although again we noted the difference in the lysosomal network in *Cln3*^Δex7/8/Δex7/8^ microglia highlighted by Lamp 2a co-staining **(Fig. 4a)**. To examine the turnover of the labelled myelin, we washed the cultured microglia and continued to incubate the cells in regular media for another 48 or 72 hours. Green-fluorescent signal was again similar in the wild-type and *Cln3*^Δex7/8/Δex7/8^ microglia at the 48-hour timepoint **(Fig. 4b, Supplementary Figure 4b)**. However, at 72 hours post-green-fluorescent-myelin washout, very little green signal was observed in the wild-type cells indicating these cells had almost completely degraded the myelin. To the contrary, significant green punctate signal was still observed in the *Cln3*^Δex7/8/Δex7/8^ microglia at 72 hours post-green-fluorescent-myelin washout, indicating these cells were not as readily able to degrade the phagocytosed green fluorescent-myelin as the wild-type microglia **(Fig. 4b,c)**.

We next studied LD formation following myelin phagocytosis. To do so, we fed the cells unlabeled myelin for 6 hours, which was followed by a wash out and regular media incubation for an additional 48 hours. LDs were then labeled with LipiDyeII. At 48 hours post-myelin washout, wild-type cells displayed nicely rounded spherical LDs that were strongly LipiDyeII positive **(Fig. 4d**; white). To the contrary, *Cln3*^Δex7/8/Δex7/8^ microglia displayed very few LDs labeled with LipiDyeII, with only a few cells harboring small LipiDyeII-positive LDs **(Fig. 4e)**. We next examined relative free cholesterol post myelin phagocytosis and washout, using Filipin dye. In contrast to LipiDyeII, Filipin staining was dramatically increased in *Cln3*^Δex7/8/Δex7/8^ microglia 48 hours post-myelin washout **(Fig. 4g)**, compared to that observed in the wild-type microglia **(Fig. 4f)**. This suggests that cholesterol derived from the phagocytosis of myelin cannot be processed and stored in LDs but instead accumulates inside CLN3-deficient microglia.

Since cholesterol was not recycled in lipid droplets but accumulated inside microglia after myelin uptake, we wanted to further investigate cholesterol metabolism in *Cln3*^Δex7/8/Δex7/8^ mouse microglia. For this, we expanded our studies, monitoring cholesterol handling kinetics in *Cln3*^Δex7/8/Δex7/8^ mouse microglia throughout the myelin uptake assay. Notably, unchallenged CLN3-deficient mice microglia prior to myelin incubation also showed significantly elevated cholesterol levels inside of lysosomes **(Fig. 4i,j,n)**. Following the 6-hour myelin incubation, both wild-type and *Cln3*^Δex7/8/Δex7/8^ microglia had significantly elevated cholesterol levels revealed by bright Filipin staining that did not significantly differ between wild-type and *Cln3*^Δex7/8/Δex7/8^ microglia, consistent with the evidence for relatively normal myelin uptake by phagocytosis by *Cln3*^Δex7/8/Δex7/8^ microglia **(Fig. 4k,n)**. However, at 48 hours post-myelin washout, as before, we observed a robust reduction in Filipin-labeled cholesterol in wild-type cells that was not evident in *Cln3*^Δex7/8/Δex7/8^ microglia, which instead still showed a significant amount of Filipin-labeled cholesterol **(Fig. 4l,n)**. Moreover, wild-type microglia showed virtually no Filipin staining by 72 hours, while the CLN3-deficient microglia still showed significant levels of Filipin-labeled cholesterol, indicating the CLN3-deficient microglia had not completely turned over the myelin-derived cholesterol at 72 hours, while the wild-type microglia had done so **(Fig. 4m,n)**.

### Autophagy inducers and cholesterol-lowering drugs rescue the myelin turnover defects in *Cln3*^Δex7/8/Δex7/8^ microglia

We have previously identified compounds that promote autophagy-lysosomal function in CLN3-deficient cells, which notably also included several FDA-approved cholesterol-lowering drugs, including statins, which are inhibitors of isoprenoid (e.g. cholesterol) biosynthesis ^52^. We next tested whether the compounds promoting autophagy-lysosomal function and/or the cholesterol-lowering drugs had any significant impact on the ability of *Cln3*^Δex7/8/Δex7/8^ microglia to efficiently degrade phagocytosed myelin and handle the myelin-associated cholesterol. For this, Lovastatin, the most effective cholesterol lowering drug from prior studies ^52^, Verapamil (an mTOR-independent autophagy inducer), and ML-SA5 [a TRPML-1 agonist that promotes TFEB activation and autophagy-lysosomal function in CLN3 disease models (Chandrachud and Cotman, unpublished data, and ^53^)] were tested. Microglia from wild-type and *Cln3*^Δex7/8/Δex7/8^ mice were fed unlabeled myelin for 6 hours, followed by a thorough washout and subsequent incubation in regular media for an additional 48 hours. In the final 24 hours of the 48-hour incubation period, we introduced ML-SA5, Verapamil, or Lovastatin (or DMSO vehicle control) to assess their potential in correcting the LD formation phenotype, using LipiDyeII to visualize LD formation. As seen in our prior experiments, CLN3-deficient microglia **(Fig. 5b)** produced less LDs than wild-type microglia **(Fig. 5a,p)**. Interestingly, all three drugs tested partially rescued the reduced LD formation **(Fig. 5c,d,e,p)**, with ML-SA5 being most efficacious. We next performed the same assay but, this time, stained with Filipin to see the effect of drug treatments on cholesterol levels. The cholesterol-reducing drug, Lovastatin, was the most effective in reducing cholesterol in *Cln3*^Δex7/8/Δex7/8^ mutant microglia **(Fig. 5j)** to a level seen in wild-type cells **(Fig. 5f,r),** but ML-SA5 also showed a significant reduction in accumulated cholesterol in the *Cln3*^Δex7/8/Δex7/8^ mutant microglia post myelin uptake/washout **(Fig. 5h,r)**. We also tested these drugs for their ability to clear the autofluorescent storage material **(Fig. 5k,l,m,n,o)**. Following a 24-hour incubation using doses previously determined to be effective in a neuronal model system ^52^, all three drugs significantly reduced the relative abundance of autofluorescent storage material, with ML-SA5 again being the most efficacious **(Fig. 5s)**.

### Reduction of myelination markers in aged *Cln3*^Δex7/8/Δex7/8^ mouse brain

The critical role of microglia in clearing myelin debris post-neuronal injury to facilitate remyelination has been well-established ^50^. Notably, thinning of the corpus callosum, a major white matter tract which primarily consists of myelinated nerve fibers, was previously reported in CLN3 disease patients and *Cln3* mice ^55,56^. Given these prior reports and the intriguing observations from our proteomics and myelin turnover studies, we hypothesize that CLN3 deficiency may impact myelination, which would lead to a reduction in myelination in the aging CLN3-deficient brain. We therefore immunostained brain sections from aged wild-type and homozygous *Cln3*^Δex7/8/Δex7/8^ mice with the myelin markers MBP and PLP. As shown in Figure 6, both markers showed significantly reduced signal in *Cln3*^Δex7/8/Δex7/8^ mice compared to wild-type mice **(Fig. 6**), lending support to our functional data that loss of CLN3 in microglia leads to altered myelin turnover in a manner that likely impacts neuronal cell health and function, particularly in the aging brain.

## Discussion

Microglia are the primary neuroimmune cells of the central nervous system (CNS) and have fundamental neuroprotective and tissue homeostatic functions through sensing the neuronal microenvironment. They protect the brain via phagocytosis and cytokine release to remove debris/pathogens and protect against infection. In response to an imbalance of microglial functions, they may become reactive and drive neuroinflammatory processes ^57^. Microglia also play a critical role in synaptic development and remodeling and in the response to neuronal injury through their ability to turnover synapse and myelin components ^54^. Notably, neuroinflammation and synaptic alterations have been reported in models of CLN3 disease and other forms of NCL ^8,26,28,30,58–61^. Consistent with findings in those prior studies, our research on microglia from a genetically accurate mouse model of CLN3 disease, *Cln3*^Δex7/8^ mice, presented in this study, also highlights evidence of elevated neuroinflammation. In addition, we have now significantly expanded upon the cell biological processes affected by loss of CLN3 function in microglia, which likely have a significant impact on the degenerative disease process observed in CLN3 disease.

Changes in the lysosomal, Golgi and mitochondria subcellular organelles have previously been documented in neurons and other cell types upon loss of CLN3 function ^6,15,41,42^. In the current study, we have similarly observed significant changes in these membrane organelles in microglia isolated from homozygous *Cln3*^Δex7/8^ mice. Moreover, consistent with the primary localization of the CLN3 protein to the endolysosome, unbiased proteomics studies revealed a dramatic impact of loss of CLN3 function on the lysosomal proteome. The two proteins with the greatest change in homozygous *Cln3*^Δex7/8/Δex7/8^ microglia, at both timepoints analyzed, were PTTG1IP and TTYH3, which were both previously identified as likely novel lysosomal membrane transporters ^45^, and in the case of PTTG1IP, induced by TFEB ^62^. TTYH3 is predicted to be a calcium-activated chloride channel ^63^. Unfortunately, there are no validated antibodies available that recognize these proteins, hampering our ability to readily follow up on these findings; however, this is of high importance for future work to better understand the potential role of these two proteins in CLN3 protein function and CLN3 disease pathophysiology. Accompanying these changes in the lysosomal proteome were changes in levels of the mTORC1 signaling complex, a major signaling hub associated with the lysosome and metabolic regulation ^44^. Prior reports of altered mTORC1 signaling in CLN3-deficiency models support these findings ^10,12,64^.

In addition to the lysosomal proteomic signature detected in CLN3-deficient microglia, other pathways were also highlighted that prompted our follow-up functional studies. These included changes in cytoskeletal proteins and cell motility/migration pathways, as well as myelin component and myelin-associated proteins and lipid-related proteins. Microglia are highly dependent on their ability to migrate towards sites of infection and injury and on their ability to phagocytose damaged or foreign debris/pathogens to neutralize these danger signals. Herein we found that CLN3-deficient microglia were unable to perform either of these processes in an efficient manner, strongly suggesting CLN3-deficent microglia cannot serve their normal neuroprotective role. In addition to reducing microglial motility, CLN3-deficiency also led to defects in myelin turnover. Uptake of myelin via phagocytosis was not specifically impaired in the microglia derived from homozygous *Cln3*^Δex7/8^ mice. However, the handling of the associated lipid was significantly altered in the CLN3-deficient microglia. Interestingly, defects in phagocytosis have also been implicated in other CLN3 deficiency model systems ^65,66^. In a recent study by Tang et al, CLN3-deficient retinal pigment epithelial cells (RPE) were less efficient in their ability to bind and take up photoreceptor outer segments (POS) ^66^. While this may seem contradictory to our findings here in which myelin uptake by CLN3-deficient microglia was normal, it may be that the different model systems, cell types and/or assays contribute to this discrepancy. Moreover, although microglia in our study displayed normal phagocytic uptake in culture dishes, it might not be the case in their natural environment given their reduced motility. In that case, CLN3-deficient microglia may not be capable of efficiently traveling to the site of injury for engulfment. Importantly, in both studies, a key role for CLN3 in the phagocytic turnover of cellular debris is demonstrated, and further investigation of phagocytic processes in CLN3 disease pathophysiology is warranted.

Despite the ability of CLN3-deficient microglia to phagocytose myelin in culture, these cells were unable to efficiently turn over the phagocytosed myelin and its associated lipids, including cholesterol, strongly implicating lipid handling as a major defective pathway upon loss of CLN3 function. Consistent with these findings, a recent study of CLN3 patient brain showed a dramatic lysosomal accumulation of cholesterol compared to age-matched controls ^67^, and CLN3 was previously implicated as a genetic regulator of LD formation ^68^. Another recent study implicated CLN3 in phospholipid catabolism. Using an elegant lysosomal enrichment technique, coupled with untargeted metabolite profiling of CLN3-deficient lysosomes, Laqtom et al identified a unique and robust lysosomal accumulation of glycerophosphodiesters (GPDs) in the absence of CLN3, together with an increase in lysophospholipids, the metabolite upstream of GPDs in the phospholipid catabolism pathway ^19^. As a result of these findings and further CLN3 complementation and transport assays in cell culture, Laqtom et al hypothesized CLN3 is required for the efflux of GPDs from the lysosome, a key process at the final stages of phospholipid catabolism ^19^.

Given the importance of lipids and lipoprotein metabolism to microglial function ^49^, these cells may be especially vulnerable to the loss of CLN3 function and its impact on phospholipid catabolism and cholesterol handling. Lipids serve as essential building blocks for cells and are an important energy source for microglia ^49^, but they also have the potential to act as pro-inflammatory stimuli. Excessive cellular accumulation of lipids can lead to toxicity and trigger apoptosis, emphasizing the delicate balance required for maintaining cellular health ^69,70^.

Although we have not specifically tested the impact of defective lipid handling and phospholipid catabolism on the survival of CLN3-deficient microglia in this study, a prior study showed a significant decrease specifically in the numbers of autofluorescent-positive microglia from *Cln3*^Δex7/8^ mice at 9 and 18 months-of-age, which was accompanied by elevated apoptosis markers, suggesting accumulation of toxic lipid species and/or an altered metabolic capacity may lead to the demise of CLN3-deficient microglia ^40^.

### Conclusion

In summary, the high level of *CLN3* expression in microglia, together with the early onset of lysosomal storage, disrupted lipid metabolism, and inflammatory signatures in *Cln3*^Δex7/8/Δex7/8^ microglia suggests that CLN3 exerts a cell-autonomous function in microglia and that microglial pathology is not solely triggered by a neurodegenerative environment. Moreover, deficient microglial function likely contributes to and/or exacerbates the dysfunction of other cell types in CLN3 disease given the importance of cell-cell interactions in the brain microenvironment. Interestingly, a comprehensive investigation into microglia from a mouse model of another neurodegenerative lysosomal storage disorder, Nieman Pick Type C disease (NPC), caused by mutations in *NPC1*, identified defects in myelin turnover and an inflammatory signature that were similar to what has been described here for homozygous *Cln3*^Δex7/8^ mouse microglia. However, in the case of *Npc1*^-/-^ mouse microglia, phagocytic uptake was enhanced and lysosomal degradation was preserved, yet myelin turnover and lipid handling, particularly of cholesterol, were impaired ^32^. Microglial lipid metabolism has also been highlighted as an important pathway in other neurodegenerative diseases including Alzheimer’s disease, multiple sclerosis, and amyotrophic lateral sclerosis (ALS) ^49,71^. Therefore, further studies to expand our understanding of CLN3 and microglial function will provide critical new insights into the role of these cells in CLN3 disease and other neurodegenerative diseases, which will be fundamental for the development of novel therapies in the future.

## Declarations

### Ethics approval and consent to participate

Studies involving use of mice underwent thorough review and approval by the Subcommittee on Research Animal Care at MGH, ensuring adherence to government regulations and guidelines for animal welfare (Protocol # 2008N000013).

## Consent for publication

Not applicable.

## Data availability

The authors declare that the data supporting the findings of this study are available within the paper and the associated supplementary information files. The mass spectrometry proteomics raw data will be made publicly available in the ProteomeXchange Consortium via the PRIDE partner repository upon publication.

## Authors’ contributions

Conceptualization: S.Y., E.B., S.T., S.L.C.; Methodology: S.Y., E.B., A.C., S.D.S. S.T., S.L.C.; Validation: S.T., S.L.C; Formal analysis: S.Y., E.B., A.C., U.C., L.M., S.A.M., S.F.L., S.T., S.L.C.; Investigation: S.Y., E.B., A.C., U.C., L.M., S.A.M., S.F.L.; Writing - original draft: S.Y., S.L.C.; Writing - review & editing: S.Y., E.B., A.C., U.C., L.M., S.D.S., S.A.M., S.F.L., S.T., S.L.C.; Supervision: S.T., S.L.C.; Project administration: S.L.C.; Funding acquisition: E.B., S.T., S.L.C.

## Acknowledgements

We thank Diane Capen for assistance in TEM studies, which were performed in the Microscopy Core of the Center for Systems Biology/Program in Membrane Biology at MGH, which is partially supported by an Inflammatory Bowel Disease Grant (DK043351) and a Boston Area Diabetes and Endocrinology Research Center (BADERC) Award (DK057521). We also thank Emily Beckett and Brigitte Demelo for their assistance in mouse husbandry and Anna Berghofer for technical assistance.

---

**Supplementary Figure 1.**
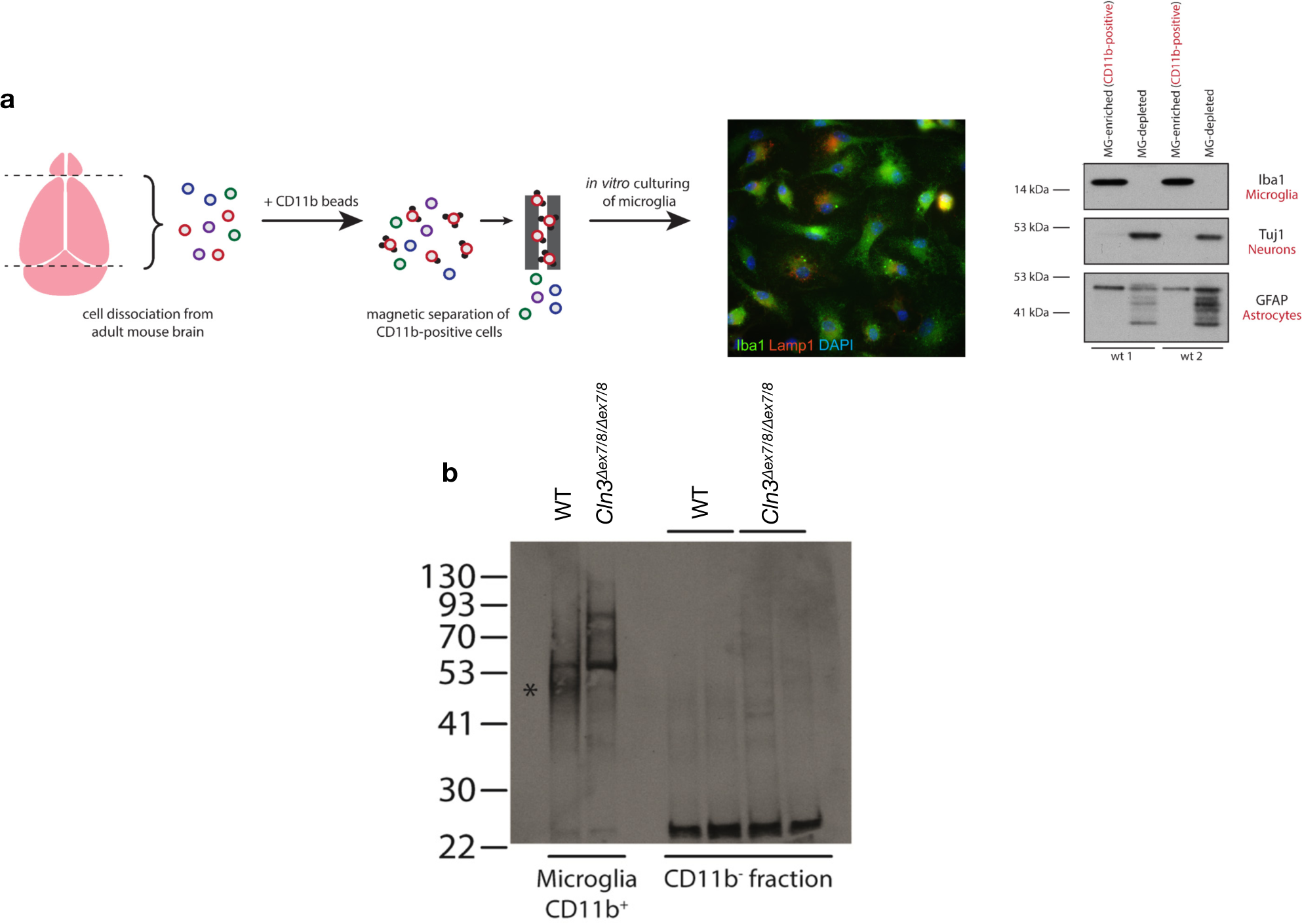
Schematic depicting MACS procedure and validation of microglial purification from mouse brain. **a)** A representative micrograph of an isolated microglial prep that was subsequently cultured and immunostained for microglial and lysosomal markers (Iba1 and Lamp1, respectively; DAPI=nuclei). Representative immunoblots for markers of microglia (Iba1), neurons (Tuj1) and astrocytes (GFAP) are shown for two independent microglia preparations from wild-type mice (‘wt 1’ and ‘wt 2’). MG=microglia. **b)** A representative immunoblot probed for the CLN3 protein is shown. 20 µg of protein lysate was loaded per lane for the microglia-enriched CD11b+ fraction and the CD11b-fraction containing primarily neurons and other types of glia. CLN3 is detectable in microglial lysates from wild-type mice (WT), but not in mutant mice (*Cln3*Δex7/8/Δex7/8) as a broad ∼50kDa band (asterisk), consistent with its highly glycosylated nature, while protein abundance in the non-microglial fraction (CD11b-fraction) is below the detection level of the antibody. Alternative higher molecular weight smears were uniquely observed in the *Cln3*Δex7/8/Δex7/8 microglial fraction; the exact nature and specificity of these bands were not further examined in the present study.

**Supplementary Figure 2.**
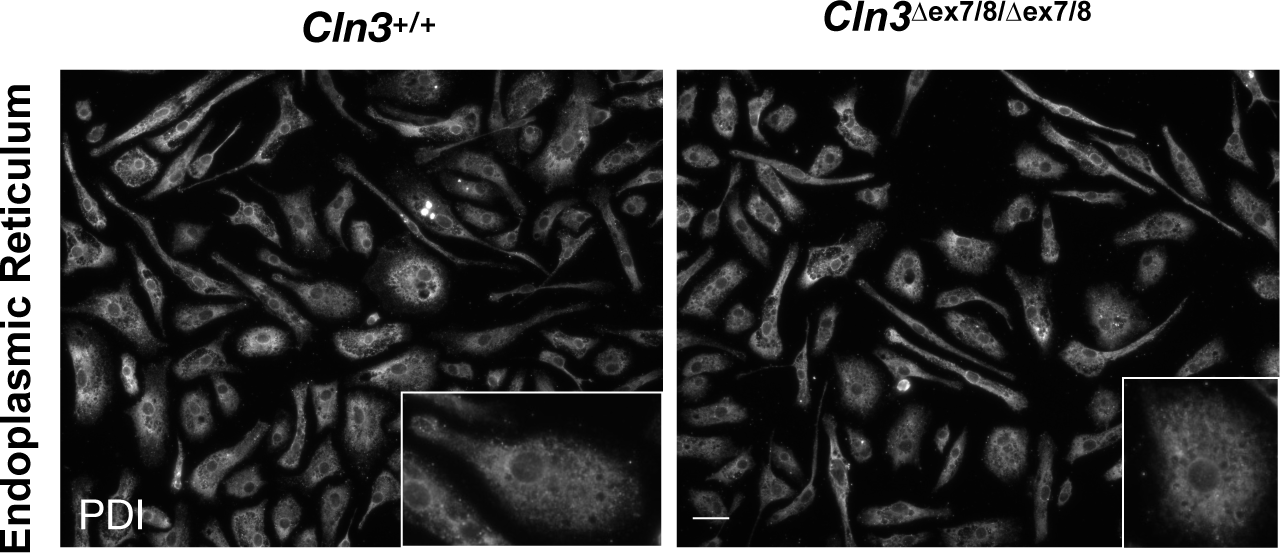
Endoplasmic reticulum (ER) morphology is normal in CLN3-deficient microglia. Representative micrograph images for ER marker PDI (white) are shown. Scale bar= 25 µm.

**Supplementary Figure 3.**
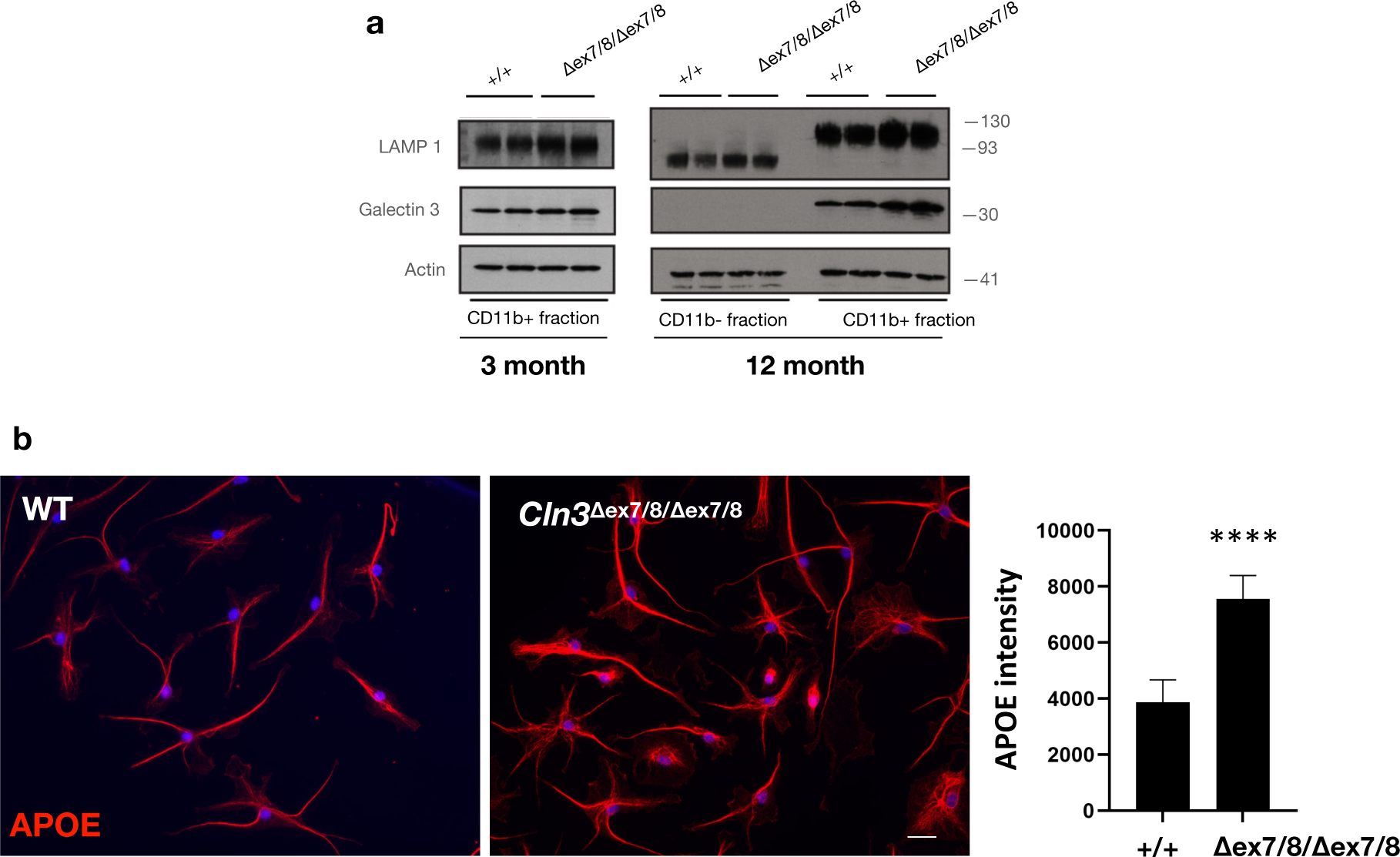
a) Increased LAMP 1 and Galectin 3 levels in CLN3-deficient microglia lysates. Immunoblots on wild-type and *Cln3*Δex7/8/Δex7/8 microglial lysates isolated from 3 and 12-month-old mice were probed with LAMP 1 and Galectin 3 antibodies. **b) Increased APOE immunostaining in CLN3-deficient microglia.** Representative micrographs of wild-type and *Cln3*Δex7/8/Δex7/8 microglial cultures immunostained using an APOE antibody are shown. The nucleus was counterstained with DAPI. The bar graph depicts the quantification of APOE intensity per cell. (N= 100 cells/ genotype, from 3 independent experiments. Data are shown as mean ± s.d.; ****P ≤ 0.0001 Student’s t-test. Scale bar= 25 µm.

**Supplementary Figure 4.**
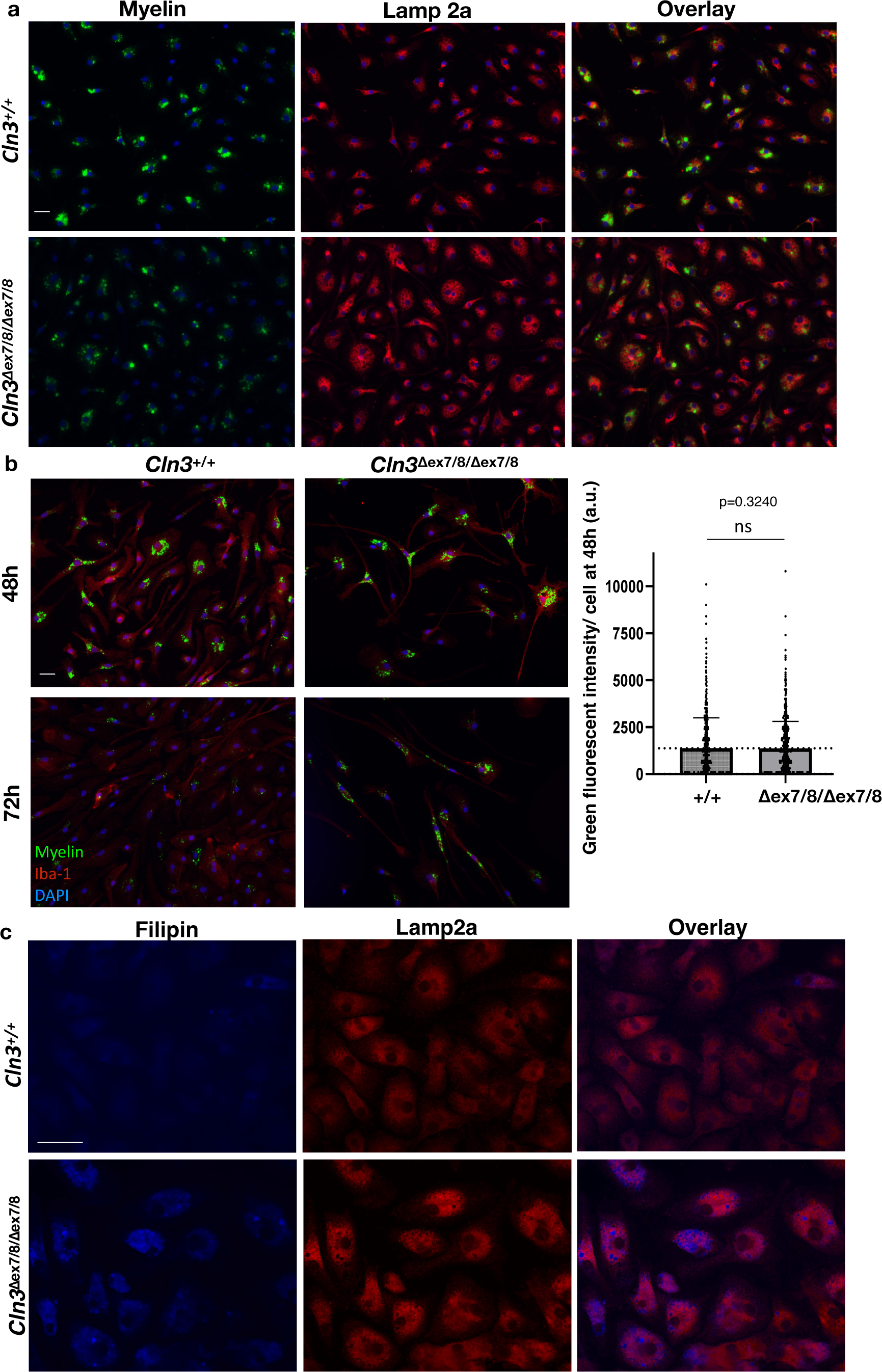
Homozygous *Cln3*Δex7/8 microglia can phagocytose myelin. **a)** Representative low-power micrographs of wild-type and *Cln3*Δex7/8/Δex7/8 microglial cultures following green fluorescent-myelin incubation for 2 hours are shown. Green fluorescent-myelin was found in Lamp2a-positive lysosomal vesicles (red). Data are representative of results from 3 independent experiments. DAPI=nuclei. Scale bar= 25 µm. **b)** Representative low-power micrographs of green-fluorescent myelin degradation at 48 and 72 hours for wild-type and homozygous *Cln3*Δex7/8 microglia. Myelin (green), Iba-1 (red), DAPI= nuclei (blue). Scale bar= 25 µm. The bar graph depicts the quantification of the green-fluorescent signal at 48 hours post-myelin uptake from three independent experiments. (N= 210 cells/genotype). Each dot represents one cell. Data are shown as mean ± s.d.; ns: non-significant, Student’s t-test. **c)** Representative micrographs from wild-type and *Cln3*Δex7/8/Δex7/8 microglial cultures co-stained with Lamp2a antibody (red) and Filipin (blue) following myelin uptake are shown. Note Filipin puncta localize to Lamp2a-positive compartments. Scale bar= 25 µm.

## Abbreviations

NCLs: Neuronal ceroid lipofuscinoses
IF: immunofluorescence
IHC: immunohistochemistry min minutes
BSA: Bovine Serum Albumin
NHS: Normal Horse Serum
PBS: Phosphate buffered saline
RT: Room temperature
O/N: Overnight
HBSS: Hank’s Balanced Salt Solution
FBS: Fetal bovine serum
NGS: Normal Goat Serum
LFQ: Label-free quantification
GOTERM_CC_FAT: gene ontology enrichment for cellular component
GOTERM_BP_FAT: gene ontology enrichment for biological process
PFA: Paraformaldehyde
LD: Lipid droplet
MACS: magnetic-activated cell sorting subunit c Subunit c of the mitochondrial ATP synthase
TEM: Transmission electron microscopy
ER: Endoplasmic reticulum
LC/MS: liquid chromatography/mass spectrometry
GO: Gene Ontology
CNS: Central nervous system
NPC: Nieman Pick Type C disease
ALS: amyotrophic lateral sclerosis

## References

1. Mole SE, Williams RE, Goebel HH. The Neuronal Ceroid Lipofuscinoses (Batten Disease) Second Edition ed. Oxford Univ. Press, Oxford; 2011.

2. Butz ES, Chandrachud U, Mole SE, Cotman SL. Moving towards a new era of genomics in the neuronal ceroid lipofuscinoses. Biochim. Biophys. Acta Mol. Basis Dis. 2020;1866,165571.

3. Mole, SE, Williams RE, Goebel HH. Correlations between genotype, ultrastructural morphology and clinical phenotype in the neuronal ceroid lipofuscinoses. Neurogenetics. 2005;6(3):107–26.

4. Radke J, Stenzel W, Goebel HH. Human NCL Neuropathology. Biochim Biophys Acta; 2015.

5. Williams RE, Mole SE. New nomenclature and classification scheme for the neuronal ceroid lipofuscinoses. Neurology. 2012;79(2):183–91.

6. Fossale E, et al. Membrane trafficking and mitochondrial abnormalities precede subunit c deposition in a cerebellar cell model of juvenile neuronal ceroid lipofuscinosis. BMC Neurosci. 2004;10;5:57.

7. Luiro K, et al. Interconnections of CLN3, Hook1 and Rab proteins link Batten disease to defects in the endocytic pathway. Human molecular genetics. 2004;13(23):3017-27.

8. Luiro K, et al. Batten disease (JNCL) is linked to disturbances in mitochondrial, cytoskeletal, and synaptic compartments. J Neurosci Res. 2006;84(5):1124–38.

9. Schultz ML, Tecedor L, Stein CS, Stamnes MA, Davidson BL. CLN3 deficient cells display defects in the ARF1-Cdc42 pathway and actin-dependent events. PLoS ONE. 2014;9(5):e96647.

10. Cao Y, et al. Autophagy is disrupted in a knock-in mouse model of juvenile neuronal ceroid lipofuscinosis. J Biol Chem. 2006;281(29):20483–93.

11. Cao Y, et al. Distinct Early Molecular Responses to Mutations Causing vLINCL and JNCL Presage ATP Synthase Subunit C Accumulation in Cerebellar Cells. PLoS ONE. 2011;6(2): e17118.

12. Vidal-Donet JM, Carcel-Trullols J, Casanova B, Aguado C, Knecht E. Alterations in ROS activity and lysosomal pH account for distinct patterns of macroautophagy in LINCL and JNCL fibroblasts. PLoS ONE. 2013;8(2):e55526.

13. Chandrachud U, et al. Unbiased Cell-based Screening in a Neuronal Cell Model of Batten Disease Highlights an Interaction between Ca2+ Homeostasis, Autophagy, and CLN3 Protein Function. J Biol Chem. 2015;290(23):14361-80.

14. Pearce DA, Nosel SA, Sherman F. Studies of pH regulation by Btn1p, the yeast homolog of human Cln3p. Molecular Genetics & Metabolism. 1999;66(4):320–3.

15. Gachet Y, Codlin S, Hyams JS, Mole SE. btn1, the Schizosaccharomyces pombe homologue of the human Batten disease gene CLN3, regulates vacuole homeostasis. J Cell Sci. 2005;1;118(Pt 23):5525-36.

16. Holopainen JM, Saarikoski J, Kinnunen PK, Jarvela I. Elevated lysosomal pH in neuronal ceroid lipofuscinoses (NCLs). European Journal of Biochemistry. 2001;268(22):5851–6.

17. Persaud-Sawin DA, McNamara JOI, Vandongen RS, Boustany R. A galactosylceramide binding domain is involved in trafficking of CLN3 from golgi to rafts via recycling endosomes. Pediatric Research. 2004;56(3):449–63.

18. Tecedor L, Stein CS, Schultz ML, Farwanah H, Sandhoff K, Davidson BL. CLN3 loss disturbs membrane microdomain properties and protein transport in brain endothelial cells. J Neurosci. 2013;33(46):18065–79.

19. Laqtom NN, et al. CLN3 is required for the clearance of glycerophosphodiesters from lysosomes. Nature. 2022;609(7929):1005-1011.

20. Rietdorf K, Coode EE, Schulz A, Wibbeler E, Bootman MD, Ostergaard JR. Cardiac pathology in neuronal ceroid lipofuscinoses (NCL): More than a mere co-morbidity. Biochim Biophys Acta Mol Basis Dis. 2020;1866(9).

21. International Batten Disease Consortium. Isolation of a novel gene underlying Batten disease, CLN3. The International Batten Disease Consortium, Cell. 1995;82(6) 949–57.

22. Eliason SL, et al. A knock-in reporter model of Batten disease. J Neurosci. 2007;12;27(37):9826-34.

23. Margraf LR, et al. Tissue expression and subcellular localization of CLN3, the Batten disease protein. Mol Genet Metab. 1999;66(4):283–9.

24. Pontikis CC, Cotman SL, MacDonald ME, Cooper JD. Thalamocortical neuron loss and localized astrocytosis in the Cln3Deltaex7/8 knock-in mouse model of Batten disease. Neurobiol Dis. 2005;20(3):823–36.

25. Burkovetskaya M, et al. Evidence for aberrant astrocyte hemichannel activity in Juvenile Neuronal Ceroid Lipofuscinosis (JNCL). PLoS One. 2014;15;9(4):e95023.

26. Parviainen L, et al. Glial cells are functionally impaired in juvenile neuronal ceroid lipofuscinosis and detrimental to neurons. Acta Neuropathol Commun. 2017;17;5(1):74.

27. Xiong J, Kielian T. Microglia in juvenile neuronal ceroid lipofuscinosis are primed toward a pro-inflammatory phenotype. J Neurochem. 2013;127(2):245–58.

28. Burkovetskaya M, Karpuk N, Kielian T. Age-dependent alterations in neuronal activity in the hippocampus and visual cortex in a mouse model of Juvenile Neuronal Ceroid Lipofuscinosis (CLN3). Neurobiol Dis. 2017;100:19–29.

29. Groh J, Ribechini E, Stadler D, Schilling T, Lutz MB, Martini R. Sialoadhesin promotes neuroinflammation-related disease progression in two mouse models of CLN disease. Glia. 2016;64(5):792–809.

30. Groh J, Berve K, Martini R. Fingolimod and Teriflunomide Attenuate Neurodegeneration in Mouse Models of Neuronal Ceroid Lipofuscinosis. Mol Ther. 2017;25(8):1889–1899.

31. Cotman SL, et al. Cln3Δex7/8 knock-in mice with the common JNCL mutation exhibit progressive neurologic disease that begins before birth. Human molecular genetics. 2002;11(22):2709–21.

32. Colombo A, et al. Loss of NPC1 enhances phagocytic uptake and impairs lipid trafficking in microglia. Nat Commun. 2021;12(1):1158.

33. Sebastian Monasor L, et al. Fibrillar Aβ triggers microglial proteome alterations and dysfunction in Alzheimer mouse models. Elife. 2020;9:e54083.

34. Tyanova S, et al. The Perseus computational platform for comprehensive analysis of (prote)omics data. Nat Methods. 2016;13(9):731–40.

35. Tusher VG, Tibshirani R, Chu G. Significance analysis of microarrays applied to the ionizing radiation response. Proc Natl Acad Sci U S A. 2001;98(9):5116–21.

36. Huang da W, Sherman BT, Lempicki RA. Systematic and integrative analysis of large gene lists using DAVID bioinformatics resources. Nat Protoc. 2009;4(1):44–57.

37. Murray MY, et al. Macrophage migration and invasion is regulated by MMP10 expression. PLoS One. 2013;8(5):e63555.

38. Panusatid C, Thangsiriskul N, Peerapittayamongkol C. Methods for mitochondrial health assessment by High Content Imaging System. MethodsX. 2022;9:101685.

39. Sun LO, et al. Spatiotemporal Control of CNS Myelination by Oligodendrocyte Programmed Cell Death through the TFEB-PUMA Axis. Cell. 2018;175(7):1811–1826.e21.

40. Burns JC, et al. Differential accumulation of storage bodies with aging defines discrete subsets of microglia in the healthy brain. Elife. 2020;9:e57495.

41. Codlin S, Mole SE. S. pombe btn1, the orthologue of the Batten disease gene CLN3, is required for vacuole protein sorting of Cpy1p and Golgi exit of Vps10p. J Cell Sci. 2009;122(Pt 8):1163–73.

42. Lojewski X, et al. Human iPSC models of neuronal ceroid lipofuscinosis capture distinct effects of TPP1 and CLN3 mutations on the endocytic pathway. Hum. Mol. Genet. 2014;23, 2005–2022.

43. Zhang Y, et al. An RNA-sequencing transcriptome and splicing database of glia, neurons, and vascular cells of the cerebral cortex. J Neurosci. 2014;34(36):11929–47.

44. Sengupta S, Peterson TR, Sabatini DM. Regulation of the mTOR complex 1 pathway by nutrients, growth factors, and stress. Mol Cell. 2010;40(2):310–22.

45. Chapel A, et al. An extended proteome map of the lysosomal membrane reveals novel potential transporters. Mol Cell Proteomics. 2013;12(6):1572–88.

46. Uusi-Rauva K, et al. Neuronal ceroid lipofuscinosis protein CLN3 interacts with motor proteins and modifies location of late endosomal compartments. Cell Mol Life Sci. 2012;69(12):2075–89.

47. Getty AL, Benedict JW, Pearce DA. A novel interaction of CLN3 with nonmuscle myosin-IIB and defects in cell motility of Cln3(-/-) cells. Exp Cell Res. 2011;317(1):51–69.

48. Bu G. Apolipoprotein E and its receptors in Alzheimer’s disease: pathways, pathogenesis and therapy. Nat Rev Neurosci 2009;10, 333–344.

49. Loving BA, Bruce KD. Lipid and Lipoprotein Metabolism in Microglia. Front Physiol. 2020;28;11:393.

50. Cantuti-Castelvetri L, et al. Defective cholesterol clearance limits remyelination in the aged central nervous system. Science. 2018;359(6376):684-688.

51. Gaultier A, et al. Lowdensity lipoprotein receptor-related protein 1 is an essential receptor for myelin phagocytosis. J Cell Sci. 2009;122:1155–1162.

52. Petcherski A, et al. An Autophagy Modifier Screen Identifies Small Molecules Capable of Reducing Autophagosome Accumulation in a Model of CLN3-Mediated Neurodegeneration. Cells. 2019;8(12):1531.

53. Wünkhaus, D., et al. TRPML1 activation ameliorates lysosomal phenotypes in CLN3 deficient retinal pigment epithelial cells. Preprint at doi: 10.1101/2023.06.21.545896 (2023).

54. Xu T, et al. The roles of microglia and astrocytes in myelin phagocytosis in the central nervous system. J Cereb Blood Flow Metab. 2023;43(3):325–340.

55. Jadav RH, et al. Magnetic Resonance Imaging in Neuronal Ceroid Lipofuscinosis and its Subtypes. The Neuroradiology Journal. 2012;25(6):755–761.

56. Palmieri M, et al. mTORC1-independent TFEB activation via Akt inhibition promotes cellular clearance in neurodegenerative storage diseases. Nat Commun. 2017;8,14338.

57. Hickman S, Izzy S, Sen P, Morsett L, El Khoury J. Microglia in neurodegeneration. Nat Neurosci. 2018;21(10):1359–1369.

58. Bosch ME, Kielian T. Neuroinflammatory paradigms in lysosomal storage diseases. Front Neurosci. 2015;9:417.

59. Song JW, et al. Lysosomal activity associated with developmental axon pruning. J Neurosci. 2008;28(36):8993–9001.

60. Llavero Hurtado M, et al. Proteomic mapping of differentially vulnerable pre-synaptic populations identifies regulators of neuronal stability in vivo. Sci Rep. 2017;7(1):12412.

61. Kielar C, et al. Molecular correlates of axonal and synaptic pathology in mouse models of Batten disease. Hum Mol Genet. 2009;18(21):4066–80.

62. Sardiello M, et al. A gene network regulating lysosomal biogenesis and function. Science. 2009;325(5939):473-7.

63. Suzuki M, Morita T, Iwamoto T. Diversity of Cl(-) channels. Cell Mol Life Sci. 2006;63(1):12–24.

64. Zhong Y, et al. Loss of CLN3, the gene mutated in juvenile neuronal ceroid lipofuscinosis, leads to metabolic impairment and autophagy induction in retinal pigment epithelium. Biochim Biophys Acta Mol Basis Dis. 2020;1866(10):165883.

65. Wavre-Shapton ST, et al. Photoreceptor phagosome processing defects and disturbed autophagy in retinal pigment epithelium of Cln3Δex1-6 mice modelling juvenile neuronal ceroid lipofuscinosis (Batten disease). Hum Mol Genet. 2015;24(24):7060–74.

66. Tang C, et al. A human model of Batten disease shows role of CLN3 in phagocytosis at the photoreceptor-RPE interface. Commun Biol. 2021;4(1):161.

67. Chen J, et al. Juvenile CLN3 disease is a lysosomal cholesterol storage disorder: similarities with Niemann-Pick type C disease. EBioMedicine. 2023;92:104628.

68. Marschallinger J, et al. Lipid-droplet-accumulating microglia represent a dysfunctional and proinflammatory state in the aging brain. Nat Neurosci. 2020;23(2):194–208.

69. Zeng X, Liu D, Huo X, Wu Y, Liu C, Sun Q. Pyroptosis in NLRP3 inflammasome-related atherosclerosis. Cell Stress 2022;6(10): 79–88.

70. Lee J, Dimitry JM, Song JH. Microglial REV-ERBα regulates inflammation and lipid droplet formation to drive tauopathy in male mice. Nat Commun 2023;14, 5197.

71. Estes RE, Lin B, Khera A, Davis MY. Lipid Metabolism Influence on Neurodegenerative Disease Progression: Is the Vehicle as Important as the Cargo? Front Mol Neurosci. 2021;14:788695.

